# Phagocytic glia are obligatory intermediates in transmission of mutant huntingtin aggregates across neuronal synapses

**DOI:** 10.1101/868448

**Authors:** Kirby M. Donnelly, Olivia R. DeLorenzo, Aprem D.A. Zaya, Gabrielle E. Pisano, Wint M. Thu, Liqun Luo, Ron R. Kopito, Margaret M. Panning Pearce

**Affiliations:** Department of Biological Sciences, University of the Sciences, Philadelphia, PA 19104, USA; Program in Neuroscience, University of the Sciences, Philadelphia, PA 19104, USA; Department of Biology, Stanford University, Stanford, CA 94305, USA; Howard Hughes Medical Institute, Stanford University, Stanford, CA 94305, USA

## Abstract

Emerging evidence supports the hypothesis that pathogenic protein aggregates associated with neurodegenerative diseases spread from cell to cell through the brain in a manner akin to infectious prions. Here, we show that mutant huntingtin (mHtt) aggregates associated with Huntington disease transfer anterogradely from presynaptic to postsynaptic neurons in the adult *Drosophila* olfactory system. Trans-synaptic transmission of mHtt aggregates is inversely correlated with neuronal activity and blocked by inhibiting caspases in presynaptic neurons, implicating synaptic dysfunction and cell death in aggregate spreading. Remarkably, mHtt aggregate transmission across synapses requires the glial scavenger receptor Draper and involves a transient visit to the glial cytoplasm, indicating that phagocytic glia act as obligatory intermediates in aggregate spreading between synaptically-connected neurons. These findings expand our understanding of phagocytic glia as double-edged players in neurodegeneration—by clearing neurotoxic protein aggregates, but also providing an opportunity for prion-like seeds to evade phagolysosomal degradation and propagate further in the brain.

## INTRODUCTION

Neurodegenerative diseases have emerged as one of the greatest healthcare challenges in our aging society, and thus a better understanding of the underlying pathological mechanisms is critical for development of more effective treatments or cures for these fatal disorders. A common molecular feature of many neurodegenerative diseases [e.g., Alzheimer disease (AD), frontotemporal dementias (FTD), Parkinson disease (PD), amyotrophic lateral sclerosis (ALS), and Huntington disease (HD)] is the misfolding of certain proteins, driving their accumulation into insoluble, amyloid aggregates (Knowles et al., 2014). Appearance of proteinaceous deposits in patient brains correlates closely with neuronal loss and clinical progression, and strategies to lower production or enhance clearance of pathological proteins in the degenerating brain have shown therapeutic promise in animal models and clinical trials (Boland et al., 2018; Li et al., 2019; Tabrizi et al., 2019).

Post-mortem histopathological analyses (Braak and Braak, 1991; Braak et al., 2003; Brettschneider et al., 2013) and *in vivo* imaging studies (Deng et al., 2004; Poudel et al., 2019) indicate that proteopathic lesions associated with neurodegeneration appear in highly-reproducible and disease-specific spatiotemporal patterns through the brain. Interestingly, these patterns largely follow neuroanatomical tracts (Ahmed et al., 2014; Mezias et al., 2017), suggesting a central role for synaptic connectivity in pathological aggregate spreading. Accumulating evidence supports the idea that intracellular aggregates formed by tau, *α*-synuclein, TDP-43, SOD1, and mutant huntingtin (mHtt) transfer from cell to cell and self-replicate by recruiting natively-folded versions of the same protein, analogous to how infectious prion protein (PrP^Sc^) templates the conformational change of soluble PrP^C^ in prion diseases (Vaquer-Alicea and Diamond, 2019). Numerous studies have pointed to roles for endocytosis and exocytosis (Asai et al., 2015; Babcock and Ganetzky, 2015; Chen et al., 2019; Holmes et al., 2013; Lee et al., 2010; Zeineddine et al., 2015), membrane permeabilization (Chen et al., 2019; Falcon et al., 2018; Flavin et al., 2017; Zeineddine et al., 2015), tunneling nanotubes (Costanzo et al., 2013; Sharma and Subramaniam, 2019), and neuronal activity (Wu et al., 2016) in entry and/or exit of pathogenic protein assemblies from cells, but the exact mechanisms by which amyloid aggregates or on-pathway intermediates cross one or more biological membranes in the highly complex central nervous system (CNS) remain an enigma.

Glia are resident immune cells of the CNS and constantly survey the brain to maintain homeostasis and respond rapidly to tissue damage or trauma. Reactive astrocytes and microglia provide a first line of defense in neurodegeneration by infiltrating sites of neuronal injury, upregulating immune-responsive genes, and phagocytosing dying neurons and other debris, including protein aggregates (Asai et al., 2015; Grathwohl et al., 2009; Wyss-Coray et al., 2003). Prolonged activation of these glial responses results in chronic inflammation, exacerbating synaptic dysfunction and neuronal loss (Hammond et al., 2018). We and others have previously shown that Draper, a *Drosophila* scavenger receptor that recognizes and phagocytoses cellular debris (Freeman, 2015), regulates the load of mHtt (Pearce et al., 2015) and A*β*_1-42_ (Ray et al., 2017) aggregate pathology in the fly CNS. Remarkably, we also found that a portion of phagocytosed neuronal mHtt aggregates gain entry into the glial cytoplasm and once there, nucleate the aggregation of normally-soluble wild-type Htt (wtHtt) proteins, suggesting that glial phagocytosis provides a path for spreading of prion-like aggregates in intact brains. Consistent with these findings, microglial ablation suppresses pathological tau transmission between synaptically-connected regions of the mouse brain (Asai et al., 2015), and PrP^Sc^ transfers from infected astrocytes to co-cultured neurons (Victoria et al., 2016). Thus, phagocytic glia may play double-edged roles in neurodegeneration, with normally neuroprotective clearance mechanisms also driving dissemination of prion-like aggregates through the brain.

A plethora of studies from the last decade have strengthened the prion-like hypothesis for neurodegenerative diseases, but we still lack a clear understanding of how pathogenic protein aggregates spread between cells in an intact CNS. In this study, we adapted our previously-described *Drosophila* HD model to investigate roles for synaptic connectivity and phagocytic glia in prion-like mHtt aggregate transmission in adult fly brains. HD is an autosomal dominant disorder caused by expansion of a CAG repeat region in exon 1 of the Htt gene, resulting in production of highly aggregation-prone mHtt proteins containing abnormally expanded polyglutamine (polyQ*≥*37) tracts (Bates et al., 2015; MacDonald et al., 1993). By contrast, wtHtt proteins containing polyQ*≤*36 tracts only aggregate upon nucleation by pre-formed Htt aggregate “seeds” (Chen et al., 2001; Preisinger et al., 1999). A growing body of evidence from cell culture (Chen et al., 2001; Costanzo et al., 2013; Holmes et al., 2013; Ren et al., 2009; Sharma and Subramaniam, 2019; Trevino et al., 2012) and *in vivo* (Ast et al., 2018; Babcock and Ganetzky, 2015; Jeon et al., 2016; Masnata et al., 2019; Pearce et al., 2015; Pecho-Vrieseling et al., 2014) models of HD supports the idea that pathogenic mHtt aggregates have prion-like properties— they transfer from cell to cell and self-replicate by nucleating the aggregation of soluble wtHtt proteins. Here, we report that mHtt aggregates formed in presynaptic olfactory receptor neuron (ORN) axons effect prion-like conversion of wtHtt proteins expressed in the cytoplasm of postsynaptic partner projection neurons (PNs) in the adult fly olfactory system. Remarkably, transfer of mHtt aggregates from presynaptic ORNs to postsynaptic PNs was abolished in Draper-deficient animals and required passage of the prion-like aggregate seeds through the cytoplasm of phagocytic glial cells. Together, these findings support the conclusion that phagocytic glia are obligatory intermediates in prion-like transmission of mHtt aggregates between synaptically-connected neurons *in vivo*, providing new insight into key roles for glia in HD pathogenesis.

## RESULTS

### Prion-like transfer of mHtt aggregates between synaptically-connected neurons in the adult fly olfactory system

Aggregates formed by N-terminal fragments of mHtt generated by aberrant splicing (e.g., exon 1; Htt_ex1_) (Sathasivam et al., 2013) or caspase cleavage (e.g., exon 1-12; Htt_ex1-12_) (Graham et al., 2006) (Fig. 1A) accumulate in HD patient brains, are highly cytotoxic, and spread between cells in culture and *in vivo* (Babcock and Ganetzky, 2015; Costanzo et al., 2013; Pearce et al., 2015; Pecho-Vrieseling et al., 2014; Ren et al., 2009). We have previously established transgenic *Drosophila* that employ binary expression systems [e.g., Gal4-UAS, QF-QUAS, or LexA-LexAop (Riabinina and Potter, 2016)] to express fluorescent protein (FP) fusions of Htt_ex1_ in non-overlapping cell populations to monitor cell-to-cell transfer of mHtt_ex1_ aggregates in intact brains (Donnelly and Pearce, 2018; Pearce et al., 2015). Our experimental approach (Fig. 1B) exploits the previously-reported finding that wtHtt_ex1_ proteins aggregate upon physically encountering mHtt_ex1_ aggregate seeds (Chen et al., 2001; Preisinger et al., 1999), such that transfer of mHtt_ex1_ aggregates from “donor” cells is reported by conversion of cytoplasmic wtHtt_ex1_ from its normally soluble, diffuse state to a punctate, aggregated state in “acceptor” cells (Fig. 1B-inset). To confirm that mHtt_ex1_ nucleates the aggregation of wtHtt_ex1_ in fly neurons, we co-expressed FP-fusions of these two proteins using pan-neuronal *elav[C155]-Gal4*. In flies expressing only mCherry-tagged mHtt_ex1_ (Htt_ex1_Q91-mCherry) pan-neuronally, aggregates were visible as discrete mCherry+ puncta throughout adult fly brains, with enrichment in neuropil regions (Fig. 1-figure supplement 1A). By contrast, GFP-tagged wtHtt_ex1_ (Htt_ex1_Q25-GFP) was expressed diffusely in the same regions of age-matched adult brains (Fig. 1-figure supplement 1B). Upon co-expression with Htt_ex1_Q91-mCherry, Htt_ex1_Q25-GFP was converted to a punctate expression pattern that almost entirely overlapped with Htt_ex1_Q91-mCherry signal (Fig. 1-figure supplement 1C and F), whereas expression patterns of neither membrane-targeted GFP (mCD8-GFP) nor soluble GFP lacking a polyQ sequence were affected by the presence of Htt_ex1_Q91-mCherry aggregates in neurons (Fig. 1-figure supplement 1D-F). Thus, in the fly CNS, Htt_ex1_Q91 aggregates induce prion-like conversion of normally-soluble Htt_ex1_Q25 via a homotypic nucleation reaction that requires the Htt_ex1_ sequence.

**Figure 1.**
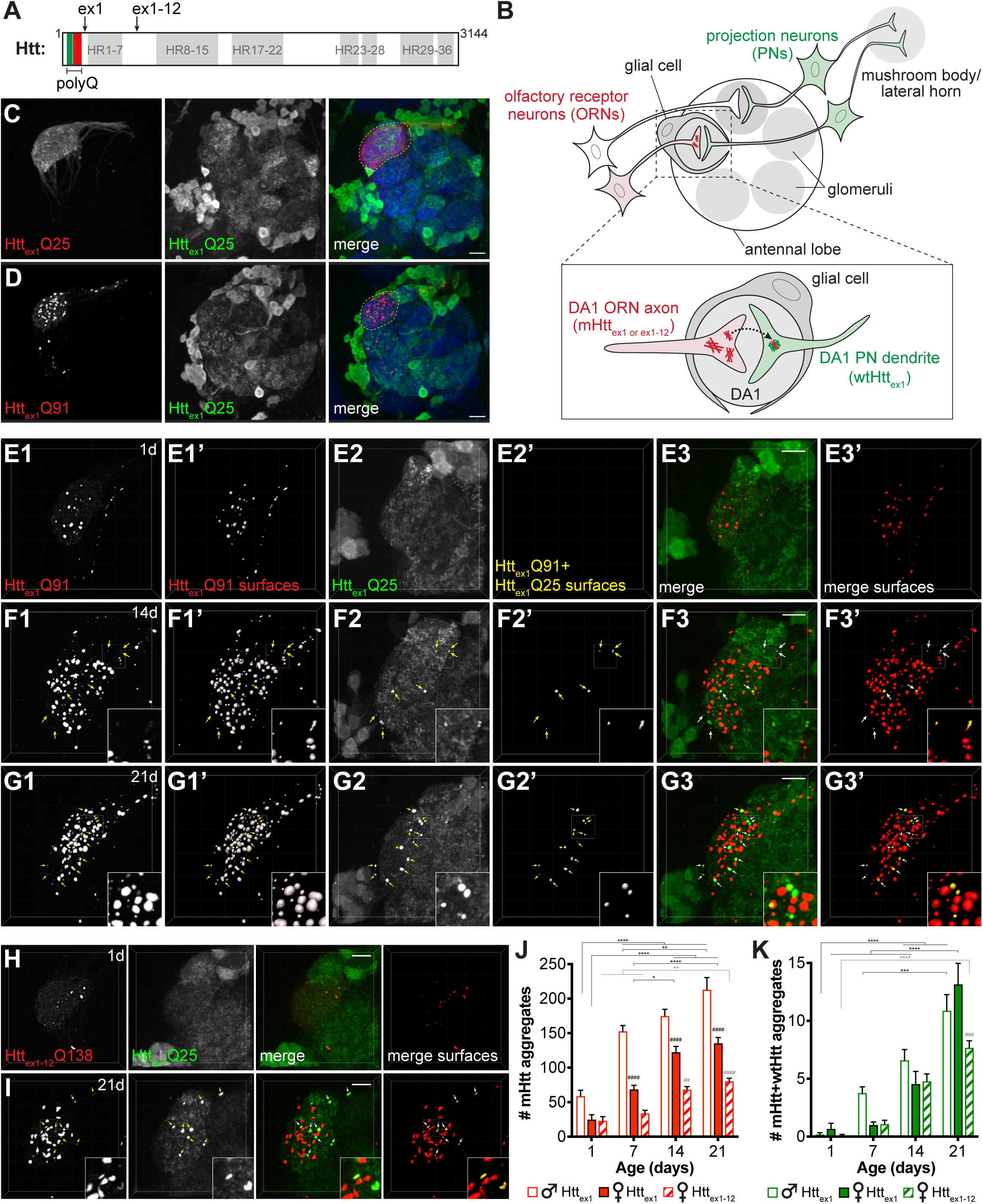
mHtt_ex1_ or mHtt_ex1-12_ aggregates formed in presynaptic ORNs induce the aggregation of wtHtt_ex1_ expressed in postsynaptic PNs. (A) Primary structure of full-length human Htt (3144 amino acids), including HEAT repeats (HR, *gray regions*) and the N-terminal variable-length polyQ region (*green/red box*), with the pathogenic threshold (∼Q37) indicated by a white dotted line. C-termini of two N-terminal mHtt fragments used in this study (Htt_ex1_ and Htt_ex1-12_) are indicated. (B) Overall experimental approach. In the fly olfactory system, ORNs synapse with PNs in discrete regions of the antennal lobe known as glomeruli (*gray circles*). PNs send axons into higher brain centers (i.e., mushroom body and/or lateral horn). Draper-expressing glial cells project processes in the antennal lobe, where they ensheath individual glomeruli. To monitor spreading of mHtt aggregates between synaptically-connected ORNs and PNs, we generated transgenic flies that express mHtt_ex1_ or mHtt_ex1-12_ fragments in DA1 ORNs and wtHtt_ex1_ in DA1 PNs. Inset: Transfer of mHtt_ex1_ or mHtt_ex1-12_ aggregates between ORNs and PNs was assessed by monitoring the solubility and colocalization of mHtt and wtHtt fluorescent signals. (C and D) Maximum intensity z-projections of antennal lobes from 7 day-old adult males expressing either Htt_ex1_Q25-mCherry (C) or Htt_ex1_Q91-mCherry (D) in DA1 ORNs using *Or67d-QF* and Htt_ex1_Q25-GFP in GH146+ PNs using *GH146-Gal4*. Raw data are shown in grayscale for individual channels and pseudocolored in merged images. Merged images include Bruchpilot immunofluorescence in blue to mark neuropil, which was used to approximate the boundaries of the DA1 glomerulus (white dotted lines). Scale bars = 20 μm. (E-G) High-magnification confocal z-stacks of DA1 glomeruli from 1 day-old (E), 14 day-old (F), and 21 day-old (G) adult males expressing Htt_ex1_Q91-mCherry in DA1 ORNs and Htt_ex1_Q25-GFP in GH146+ PNs. Boxed regions in (F and G) are shown at higher magnification in insets. Raw data are shown in grayscale in individual channels (Htt_ex1_Q91: E1, F1, G1; Htt_ex1_Q25: E2, F2, G2) and pseudocolored in merged images (E3, F3, G3). mCherry+ “Htt_ex1_Q91 surfaces” (E1’, F1’, G1’) and “Htt_ex1_Q91+Htt_ex1_Q25 surfaces” (E2’, F2’, G2’) identified by semi-automated image segmentation are shown adjacent to raw data and pseudocolored red and yellow, respectively, in the “merged surfaces” images (E3’, F3’, G3’). Arrows (*yellow* on grayscale images, *white* on merged images) indicate Htt_ex1_Q91+Htt_ex1_Q25 surfaces. Scale bars = 10 μm. (H and I) Confocal z-stacks from 1 day-old (H) and 21 day-old (I) adult females expressing RFP-Htt_ex1-12_Q138 in DA1 ORNs and Htt_ex1_Q25-GFP in GH146+ PNs. Boxed region in (I) is shown at higher magnification in insets. RFP+ surfaces identified by semi-automated image segmentation are shown in the last column, with Htt_ex1-12_Q138-only surfaces in red and Htt_ex1-12_Q138+Htt_ex1_Q25 surfaces in yellow. Scale bars = 10 μm. (J) and (K) Numbers of Htt_ex1_Q91 or Htt_ex1-12_Q138 (”mHtt”) surfaces (J) and Htt_ex1_Q91+Htt_ex1_Q25 or Htt_ex1-12_Q138+Htt_ex1_Q25 (“mHtt+wtHtt”) surfaces (K) identified in adult males (*open bars*) or females (*solid bars*) expressing Htt_ex1_Q91-mCherry in DA1 ORNs or adult females expressing RFP-Htt_ex1-12_Q138 in DA1 ORNs (*striped bars*) at the indicated ages. Data are shown as mean ± SEM; *p < 0.05, **p < 0.01, ***p < 0.001, or ****p < 0.0001 by two-way ANOVA followed by Tukey’s multiple comparisons tests. *s indicate statistical significance comparing flies of the same genotype and sex at different ages (black *s compare males or females expressing Htt_ex1_Q91, and gray *s compare females expressing Htt_ex1-12_Q138 over time). #s indicate statistical significance comparing different genotypes at the same age (black #s compare males vs females expressing Htt_ex1_Q91, and gray #s compare females expressing Htt_ex1_Q91 vs. females expressing Htt_ex1-12_Q138).

To examine trans-synaptic prion-like transfer of mHtt_ex1_ aggregates, we coupled QF-driven expression of Htt_ex1_Q91-mCherry with Gal4-driven expression of Htt_ex1_Q25-GFP in neuronal cell populations that make well-defined synaptic connections in the adult fly olfactory system (Fig. 1B-D). Htt_ex1_Q91-mCherry was expressed using *Or67d-QF* in ∼40 presynaptic ORNs (“DA1 ORNs”) that project axons from the antenna into the central brain, where they form synaptic connections with dendrites of ∼7 partner PNs (“DA1 PNs”) in the DA1 glomerulus of the antennal lobe (Fig. 1B-inset, C, and D) (Jefferis et al., 2001). In these same animals, Htt_ex1_Q25-GFP was expressed in ∼60% of PNs using *GH146-Gal4*, which labels lateral and ventral DA1 PNs in addition to other PN types (Fig. 1C and D) (Marin et al., 2002). We did not detect expression of Htt_ex1_Q91-mCherry in DA1 ORNs until ∼24 hr before eclosion, consistent with activation of adult olfactory receptor gene expression during late pupal development (Clyne et al., 1999), whereas Htt_ex1_Q25-GFP was expressed in PNs via *GH146-Gal4* earlier in development (Stocker et al., 1997). This genetic approach therefore enables us to monitor prion-like transfer of mHtt aggregates between post-mitotic, synaptically-connected DA1 ORNs and PNs in the adult fly brain.

Formation and prion-like transfer of mHtt_ex1_ aggregates across DA1 ORN-PN synapses was examined by monitoring the solubility of Htt_ex1_Q91-mCherry and Htt_ex1_Q25-GFP proteins in or near the DA1 glomerulus (Fig. 1B-inset and E-G). Whereas Htt_ex1_Q25-mCherry was expressed diffusely in DA1 ORN axons (Fig. 1C), aggregated Htt_ex1_Q91-mCherry was visible as discrete puncta almost entirely restricted to DA1 axons and axon termini in the DA1 region of the antennal lobe (Fig. 1D and E1-G1). We used semi-automated 3D segmentation and reconstruction of high-magnification confocal z-stacks (Fig. 1-figure supplement 2A1 and Video 1) to quantify Htt_ex1_Q91 aggregate formation in the DA1 glomerulus over time. Htt_ex1_Q91 aggregates first appeared in pharate adults and increased in number as the flies aged (Fig. 1E1’-G1’ and 1J). Numbers of Htt_ex1_Q91 aggregates in males exceeded those in females at each time point (Fig. 1J), consistent with known sexual dimorphism in DA1 glomerular volume (Stockinger et al., 2005). In these same brains, Htt_ex1_Q25-GFP was expressed diffusely throughout GH146+ PN cell bodies and processes in young adults (Fig. 1D and E2), but bright Htt_ex1_Q25-GFP puncta began to appear and accumulate in the DA1 glomerulus as the flies aged (Fig. 1F2 and G2, *arrows*). Because these puncta could be difficult to distinguish from surrounding non-aggregated Htt_ex1_Q25-GFP signal, and GFP+ puncta representing normal dendritic architecture and/or intracellular vesicles in the secretory pathway were visible in GH146+ PNs expressing mCD8-GFP (Fig. 1-figure supplement 2B2 and Fig. 1-figure supplement 3B), we defined Htt_ex1_Q25 aggregates as GFP+ puncta that colocalized with Htt_ex1_Q91 aggregates (Fig. 1-figure supplement 2A1 and C1-7, and Video 1). This approach reported similar results to manual quantification of the same data in 2D confocal slices (Fig. 1-figure supplement 2C1-7 and D), and identical data were obtained when Htt_ex1_Q25-GFP+ segmented surfaces were filtered for colocalization with Htt_ex1_Q91-mCherry+ puncta (Fig. 1-figure supplement 2A2 and D). By contrast, Htt_ex1_Q91 aggregates in DA1 ORNs did not colocalize with mCD8-GFP expressed in GH146+ PNs in control animals regardless of whether mCherry+ or GFP+ surfaces were initially segmented (Fig. 1-figure supplement 2B1-2). Thus, our semi-automated approach to identify “Htt_ex1_Q91+Htt_ex1_Q25” aggregates specifically reports non-cell autonomous conversion of postsynaptic wtHtt_ex1_ by presynaptic mHtt_ex1_ seeds (Fig. 1B-inset).

Numbers of Htt_ex1_Q91+Htt_ex1_Q25 aggregates increased as flies expressing Htt_ex1_Q91-mCherry in DA1 ORNs and Htt_ex1_Q25-GFP in GH146+ PNs aged from 1 to 21 days old (Fig. 1E2’-G2’ and 1J). Htt_ex1_Q91 aggregates outnumbered Htt_ex1_Q91+Htt_ex1_Q25 aggregates at each time point tested (Fig. 1J and K), likely reflecting a higher rate of aggregate formation in “donor” ORNs expressing polyQ-expanded mHtt_ex1_ than in “acceptor” PNs, where wtHtt_ex1_ proteins must be nucleated by mHtt_ex1_ seeds originating in other cells. In control experiments, soluble Htt_ex1_Q25-mCherry expressed in DA1 ORNs did not colocalize with Htt_ex1_Q25-GFP expressed in PNs (Fig. 1-figure supplement 3A, E, and F), and Htt_ex1_Q91 aggregates in DA1 ORNs did not nucleate membrane-bound mCD8-GFP in PNs (Fig. 1-figure supplement 3B, E, and F). Htt_ex1_Q91+Htt_ex1_Q25 aggregates still formed when the Gal4 inhibitor Gal80 was expressed in ORNs (Fig. 1-figure supplement 3C, E, and F) or when the QF repressor QS was expressed in PNs (Fig. 1-figure supplement 3D-F), arguing strongly against co-expression of Htt_ex1_Q91-mCherry and Htt_ex1_Q25-GFP using the highly-specific *Or67d-QF* and *GH146-Gal4* drivers. Together, these findings indicate that mHtt_ex1_ aggregates formed in presynaptic ORNs induce non-cell autonomous, homotypic aggregation of wtHtt_ex1_ expressed in the cytoplasm of postsynaptic PNs.

Htt_ex1_Q91+Htt_ex1_Q25 aggregates were not detected in non-synaptically-connected GH146+ PNs, including in glomeruli directly adjacent to where DA1 ORNs terminate, suggesting that prion-like conversion of Htt_ex1_Q25 in PNs by Htt_ex1_Q91 aggregates in ORNs requires synaptic connectivity. To examine whether Htt_ex1_Q91 aggregates can also spread retrogradely across ORN-PN synapses, we expressed Htt_ex1_Q91-mCherry in PNs using *GH146-QF* and Htt_ex1_Q25-GFP in all ORNs using *pebbled-Gal4*. While many Htt_ex1_Q91 aggregates were visible in PN soma, dendrites, and axons in adult brains (Fig. 1-figure supplement 4A-C), we did not observe colocalization of Htt_ex1_Q25-GFP and Htt_ex1_Q91-mCherry puncta within the antennal lobe neuropil in 1, 7, and 14 day-old adults (Fig. 1-figure supplement 4A, B, and D), suggesting that Htt_ex1_Q91 aggregate spreading across ORN-PN synapses is restricted to or much more efficient in the anterograde direction. We also found that aggregates formed by mHtt_ex1-12_ caspase-6 cleavage products (RFP-Htt_ex1-12_Q138) (Fig. 1A) (Graham et al., 2006), which have previously been shown to spread from ORN axons to more distant (non-PN) neurons in the fly CNS (Babcock and Ganetzky, 2015), also transfer anterogradely across DA1 ORN-PN synapses (Fig. 1H and I). Together, these findings indicate that multiple pathogenic N-terminal mHtt fragments share the ability to spread trans-synaptically in *Drosophila* brains, and the sequences required for prion-like conversion of wtHtt reside within Htt_ex1_.

### wtHtt_ex1_ aggregates in PNs are seeded by smaller mHtt_ex1_ aggregates originating in ORNs

Higher magnification examination of DA1 glomeruli in flies expressing Htt_ex1_Q91-mCherry (Fig. 2A and B) or RFP-Htt_ex1-12_Q138 (Fig. 2C and D) in DA1 ORNs and Htt_ex1_Q25-GFP in GH146+ PNs revealed that most (>85%) Htt_ex1_Q25-GFP puncta in DA1 PNs colocalized with Htt_ex1_Q91-mCherry aggregates. Segmentation in both the red (Fig 2A1’-D1’) and green (Fig. 2A2’-D2’) channels demonstrated that GFP+ surfaces entirely surrounded associated mCherry+ or RFP+ surfaces in these colocalized aggregates (Fig 2A3’-D3’). These data suggest that aggregated Htt_ex1_Q91 or Htt_ex1-12_Q138 proteins form the “core” of induced Htt_ex1_Q25 aggregates and supports our hypothesis that wtHtt_ex1_ solubility in PNs is altered upon direct physical interaction with pre-formed mHtt seeds. To further examine molecular interactions between Htt_ex1_Q91 and Htt_ex1_Q25 proteins, we measured fluorescence resonance energy transfer (FRET) in colocalized aggregates. Remarkably, positive FRET signal was detected for all Htt_ex1_Q91+Htt_ex1_Q25 aggregates analyzed by this method (Fig. 2E), indicating that the FP tags fused to each of these proteins were in close molecular proximity (<10 nm apart), consistent with direct physical contact.

**Figure 2.**
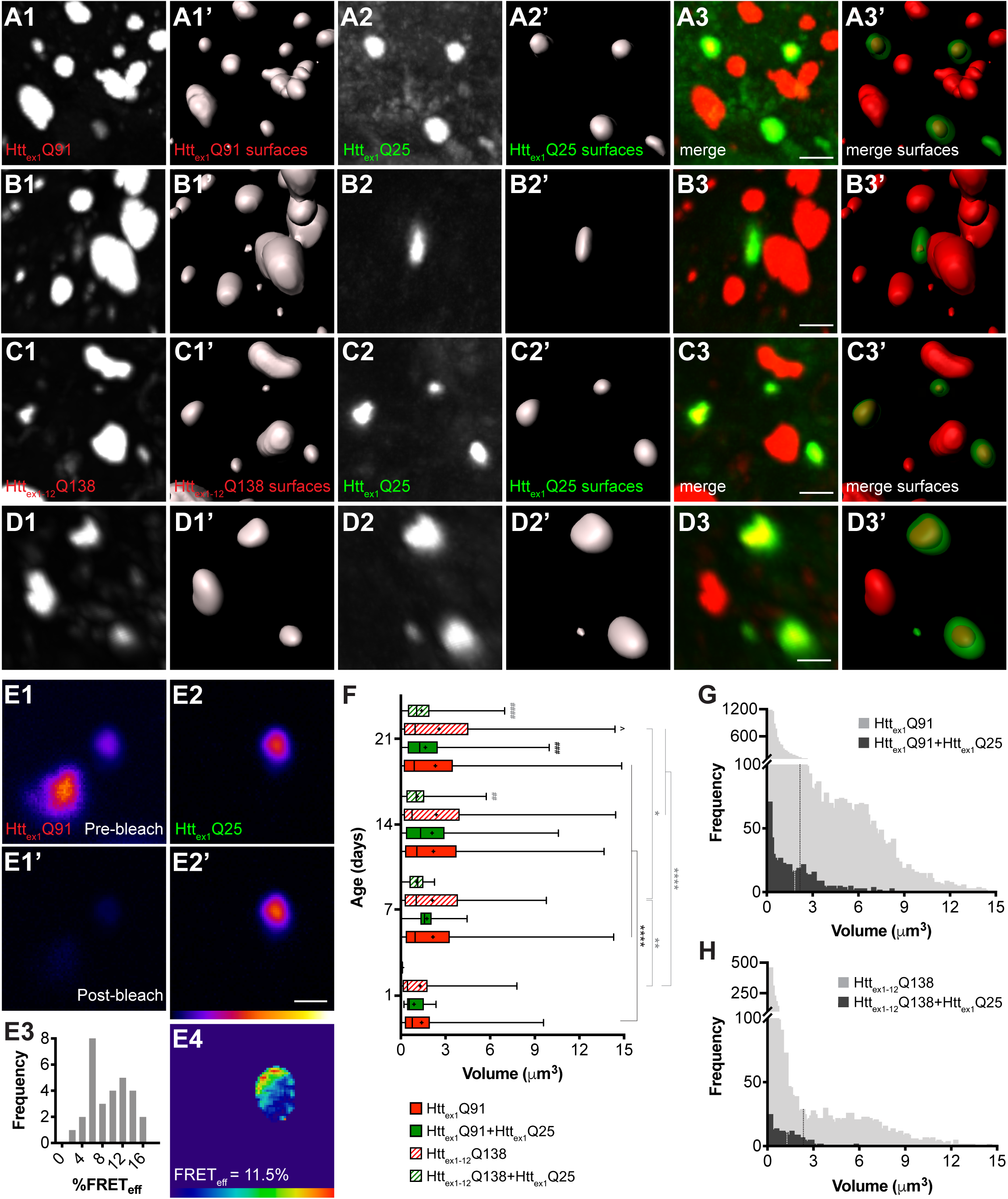
wtHtt_ex1_ aggregates in postsynaptic PNs are nucleated by mHtt_ex1_ or mHtt_ex1-12_ aggregates from presynaptic ORNs. (A-D) High-magnification confocal z-stacks of DA1 glomeruli from adult flies expressing Htt_ex1_Q91-mCherry (A and B) or RFP-Htt_ex1-12_Q138 (C and D) in DA1 ORNs and Htt_ex1_Q25-GFP in GH146+ PNs. Raw data (A1-3, B1-3, C1-3, D1-3) are shown adjacent to surfaces identified by 3D segmentation of the red (A1’, B1’, C1’, D1’) or green (A2’, B2’, C2’, D2’) channels. Htt_ex1_Q25-GFP surfaces are shown at 50% transparency in “merged surfaces” images (A3’, B3’, C3’, D3’) for visibility of co-localized Htt_ex1_Q91-mCherry or RFP-Htt_ex1-12_Q138 surfaces. Scale bars = 1 μm. (E1-4) A single confocal slice through the center of a Htt_ex1_Q91+Htt_ex1_Q25 aggregate before (E1, E2) and after (E1’, E2’) mCherry acceptor photobleaching. Data are shown as a heat map to highlight changes in fluorescence intensities after photobleaching. Scale bar = 1 μm. FRET efficiency (FRET_eff_) for this aggregate is shown in (E4), and average FRET_eff_ values for all Htt_ex1_Q91+Htt_ex1_Q25 aggregates tested are shown in (E3). (F) Volumes of Htt_ex1_Q91 (*solid red boxes*), Htt_ex1_Q91+Htt_ex1_Q25 (*solid green boxes*), Htt_ex1-12_Q138 (*striped red boxes*), and Htt_ex1-12_Q138+Htt_ex1_Q25 (*striped green boxes*) aggregates identified in the DA1 glomerulus at the indicated ages. Box widths indicate interquartile ranges, vertical lines inside each box indicate medians, whiskers indicate minimums/maximums, and “+”s indicate means for each data set. *p < 0.05, **p < 0.01, ***p < 0.001, ****p < 0.0001 by one-way ANOVA followed by Tukey’s multiple comparisons test. Statistical significance is indicated by “*”s when comparing the same aggregate sub-population at different ages (Htt_ex1_Q91 surfaces in *black* and Htt_ex1-12_Q138 surfaces in *gray*), by “#”s when comparing Htt_ex1_Q91 vs Htt_ex1_Q91+Htt_ex1_Q25 (*black*) or Htt_ex1-12_Q138 vs Htt_ex1-12_Q138+Htt_ex1_Q25 (*gray*) aggregates at the same ages, and by “^”s when comparing Htt_ex1_Q91 vs Htt_ex1-12_Q138 aggregates at the same ages. (G and H) Distribution of volumes for (G) Htt_ex1_Q91 (*light gray bars*) and Htt_ex1_Q91+Htt_ex1_Q25 (*dark gray bars*) or (H) Htt_ex1-12_Q138 (*light gray bars*) and Htt_ex1-12_Q138+Htt_ex1_Q25 (*dark gray bars*) aggregates, combined from 7, 14, and 21 day-old flies. Mean volume of Htt_ex1_Q91 or Htt_ex1-12_Q138 aggregates and Htt_ex1_Q91+Htt_ex1_Q25 or Htt_ex1-12_Q138+Htt_ex1_Q25 aggregates are indicated by black and white dotted lines, respectively, on each histogram.

Comparison of volumes for all segmented Htt_ex1_Q91, Htt_ex1-12_Q138, Htt_ex1_Q91+Htt_ex1_Q25, and Htt_ex1-12_Q138+Htt_ex1_Q25 aggregates revealed that Htt_ex1_Q91 and Htt_ex1-12_Q138 aggregates increased in size as the flies aged, most substantially during the first week of adulthood (Fig. 2F). Interestingly, the mean volume of Htt_ex1_Q91 or Htt_ex1-12_Q138 surfaces that colocalized with Htt_ex1_Q25 aggregates (herein referred to as “seeding-competent” mHtt aggregates) was less than the mean volume of all Htt_ex1_Q91 or Htt_ex1-12_Q138 surfaces, and these differences were statistically significant in older animals. When aggregate volumes were analyzed across all time points, it became apparent that seeding-competent Htt_ex1_Q91 or Htt_ex1-12_Q138 aggregates clustered in a significantly smaller-sized subpopulation (range = 0.1–3.5 μm^3^; mean = 1.845 ± 0.068 μm^3^ for Htt_ex1_Q91 aggregates, mean = 1.293 ± 0.071 μm^3^ for Htt_ex1- 12_Q138 aggregates), whereas the entire population of Htt_ex1_Q91 or Htt_ex1-12_Q138 aggregates was more heterogeneously-sized (range = 0.1–15 μm^3^; mean = 2.19 ± 0.024 μm^3^ for Htt_ex1_Q91 aggregates, mean = 2.40 ± 0.048 μm^3^ for Htt_ex1-12_Q138 aggregates) (Fig. 2G and H). We previously reported that Htt_ex1_Q25 aggregates seeded in the cytoplasm of glial cells colocalized with a similarly smaller-sized subpopulation of seeding-competent Htt_ex1_Q91 aggregates from DA1 ORNs (Pearce et al., 2015), and smaller mHtt_ex1_ aggregates are associated with increased seeding-propensity and neurotoxicity in other HD models (Ast et al., 2018; Chen et al., 2001). Taken together, these findings strongly suggest that mHtt_ex1_ or mHtt_ex1-12_ aggregates formed in presynaptic ORNs effect prion-like conversion of wtHtt_ex1_ in the cytoplasm of postsynaptic PNs, and that mHtt aggregate transmissibility in the fly CNS is correlated with smaller aggregate size.

### mHtt_ex1_ aggregate transfer is enhanced across silenced DA1 ORN-PN synapses

Endocytosis, exocytosis, and neuronal activity have been previously implicated in neuron-to-neuron spreading of mHtt and other pathogenic aggregates (Babcock and Ganetzky, 2015; Pecho-Vrieseling et al., 2014; Wu et al., 2016), but it is not known how these processes contribute to aggregate transfer across endogenous synapses *in vivo*. To examine a role for synaptic activity in DA1 ORN-to-PN transfer of mHtt_ex1_ aggregates, we used well-established fly genetic tools that block fission or fusion of synaptic vesicles at the presynaptic membrane to impair neurotransmission. First, we used *shibire^ts1^* (*shi^ts1^*) (Kosaka and Ikeda, 1983), a temperature-sensitive mutant of the GTPase Shibire/dynamin that blocks endocytic recycling of synaptic vesicles in flies raised at the restrictive temperature. Co-expression of shi^ts1^ with Htt_ex1_Q91-mCherry in DA1 ORNs in flies shifted from the permissive temperature (18°C) to the restrictive temperature (31°C) in adulthood had no effect or slightly decreased numbers of Htt_ex1_Q91 aggregates in the DA1 glomerulus (Fig. 3A-C), but, surprisingly, strongly enhanced formation of seeded Htt_ex1_Q25 aggregates in DA1 PN dendrites compared with control flies expressing LacZ (Fig. 3A, B, and D). Likewise, co-expression of Htt_ex1_Q91- mCherry with tetanus toxin light chain (TeTxLC), which inhibits SNARE-mediated fusion of synaptic vesicles with presynaptic membranes (Sweeney et al., 1995), strongly increased numbers of seeded Htt_ex1_Q25 aggregates in the DA1 glomerulus (Fig. 3E, F, and H). At some time points, TeTxLC co-expression led to increased numbers of Htt_ex1_Q91 aggregates compared with control animals (Fig. 3E-G); however, there appeared to be no correlation between numbers of Htt_ex1_Q91 and Htt_ex1_Q91+Htt_ex1_Q25 aggregates in the DA1 glomerulus over time, so the increased numbers of seeded Htt_ex1_Q25 aggregates were unlikely to be simply due to abundance of presynaptic Htt_ex1_Q91 seeds. To manipulate neuronal activity by an alternative approach, we co-expressed the heat-activated *Drosophila* transient-receptor potential A (dTrpA) channel (Hamada et al., 2008) with Htt_ex1_Q91- mCherry to thermogenetically stimulate DA1 ORNs. In adult flies shifted from ∼21°C to 31°C upon eclosion, dTrpA-mediated activation of Htt_ex1_Q91-mCherry-expressing DA1 ORNs slightly increased Htt_ex1_Q91 aggregate numbers, but decreased formation of seeded Htt_ex1_Q25 aggregates in the DA1 glomerulus compared with control flies expressing LacZ (Fig. 3I-L). Together, these results indicate that prion-like transmission of Htt_ex1_Q91 aggregates from ORNs to PNs is inversely correlated with presynaptic ORN activity. These findings suggest that aggregate transfer could be enhanced across dysfunctional synapses, which are an early pathological finding in HD and other neurodegenerative diseases.

**Figure 3.**
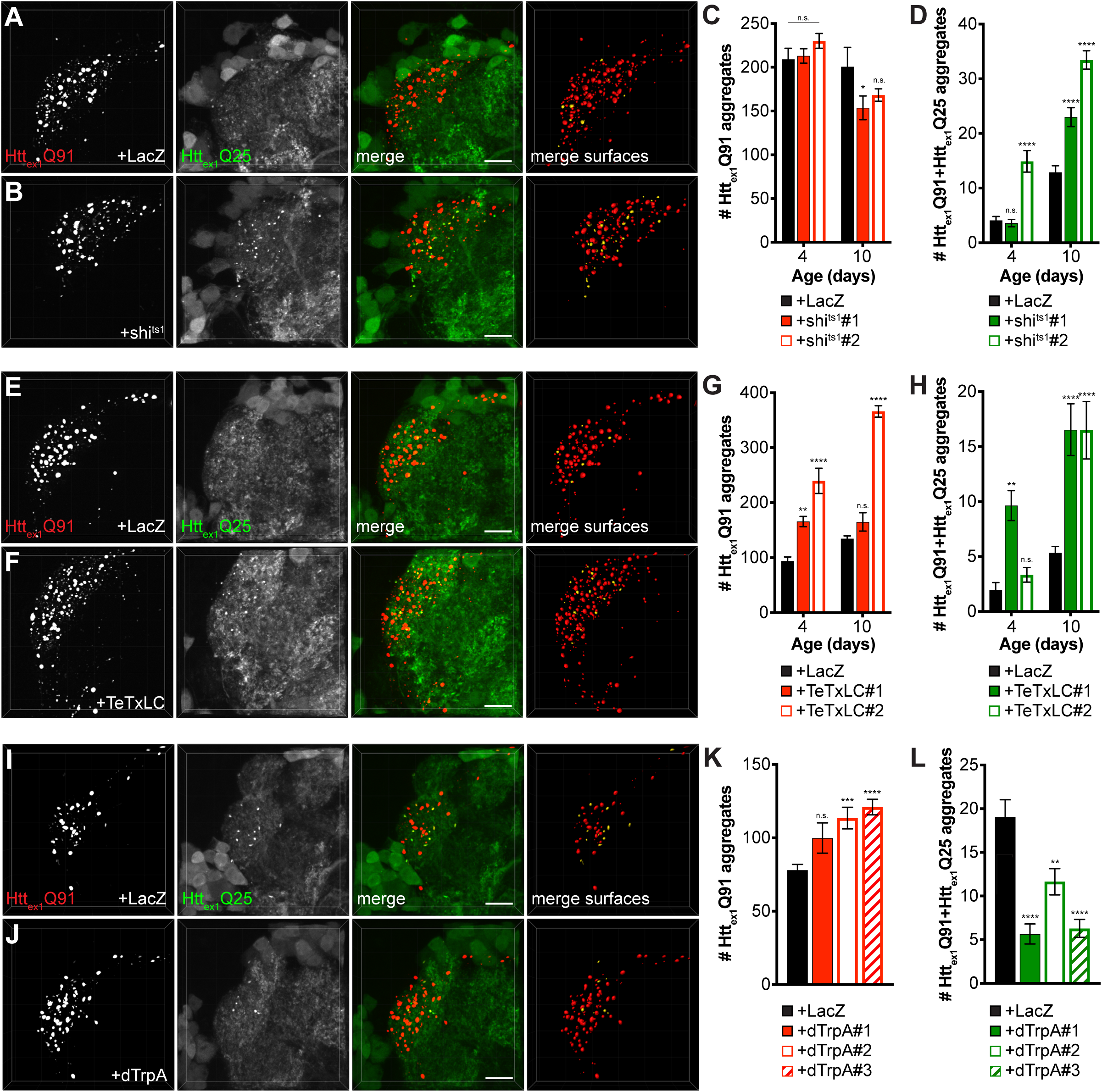
mHtt_ex1_ aggregate transfer from ORNs to synaptically-connected PNs is inversely correlated with presynaptic activity. (A, B, E, F, I, and J) Confocal z-stacks of DA1 glomeruli from 10 day-old males (A and B, and E and F) or 7 day-old females (I and J) co-expressing Htt_ex1_Q91-mCherry with either LacZ (A, E, and I), shi^ts1^ (B), TeTxLC (F), or dTrpA (J) in DA1 ORNs and Htt_ex1_Q25-GFP in GH146+ PNs. In (A-B), flies were raised at the permissive temperature (18°C) and shifted to the restrictive temperature (31°C) upon eclosion, and in (I-J), flies were raised at room temperature (∼21°C) and shifted to 31°C upon eclosion. mCherry+ surfaces identified by semi-automated image segmentation are shown in the last panels, with Htt_ex1_Q91-only surfaces in red and Htt_ex1_Q91+Htt_ex1_Q25 surfaces in yellow. Scale bars = 10 μm. (C and D, G and H, and K and L) Quantification of Htt_ex1_Q91 (C, G, and K) and Htt_ex1_Q91+Htt_ex1_Q25 (D, H, and L) aggregates identified in DA1 glomeruli from adult males of the indicated ages co-expressing Htt_ex1_Q91-mCherry with LacZ or shi^ts1^ using two independent *QUAS-shi^ts1^* lines in DA1 ORNs and Htt_ex1_Q25-GFP in GH146+ PNs (C and D), adult males of the indicated ages co-expressing Htt_ex1_Q91-mCherry with LacZ or TeTxLC using two independent *QUAS-TeTxLC* lines in DA1 ORNs and Htt_ex1_Q25-GFP in GH146+ PNs (G and H), and 7 day-old females expressing Htt_ex1_Q91-mCherry with either LacZ or dTrpA using three independent *QUAS-dTrpA* lines in DA1 ORNs and Htt_ex1_Q25-GFP in GH146+ PNs (K and L). Data are shown as mean ± SEM; *p < 0.05, **p < 0.01, ***p < 0.001, ****p < 0.0001, n.s. = not significant by one- or two-way ANOVA with Tukey’s multiple comparisons test comparing shi^ts1^-, TeTxLC-, or dTrpA-expressing flies to their respective controls expressing LacZ.

Our results using shi^ts1^ and TeTxLC to block DA1 ORN activity contrast with previous reports showing that spreading of Htt_ex1-12_Q138 aggregates was inhibited from endocytosis- or exocytosis-impaired ORNs to non-synaptically-connected neurons (Babcock and Ganetzky, 2015) or that botulinum toxin inhibited spreading of mHtt_ex1_ from R6/2 mouse brain slices to functionally-connected human neurons (Pecho-Vrieseling et al., 2014). This discrepancy could be due to construct- or cell type-specific effects or possibly different mechanisms regulating synaptic or non-synaptic aggregate transmission in the brain. Thus, we wondered whether blocking Shibire-mediated endocytosis in ORNs might create a more favorable environment for aggregate transfer across endogenous synapses in our HD model. To test this, we first quantified Htt_ex1_Q91-mCherry-expressing DA1 ORN axonal surfaces using mCD8-GFP, a tool widely used to label neuronal cell bodies and processes (Lee and Luo, 1999; Mosca and Luo, 2014) and to quantify neuron or neurite abundance (Burr et al., 2014; MacDonald et al., 2006) in fly brains. Segmentation and 3D reconstruction of mCD8-GFP+ DA1 ORN surfaces revealed that fluorescence intensity of and volume occupied by DA1 ORN axons were increased ∼2-fold in shi^ts1^-expressing flies at the restrictive temperature compared with controls expressing LacZ (Fig. 4A-C). This effect appeared to be specific since Htt_ex1_Q91 aggregate abundance in DA1 ORNs was not increased by shi^ts^ co-expression (Fig. 3C), and may reflect a homeostatic compensatory response to endocytic blockade (Davis, 2013; Dickman et al., 2006) that could create additional exit sites for Htt_ex1_Q91 aggregates.

**Figure 4.**
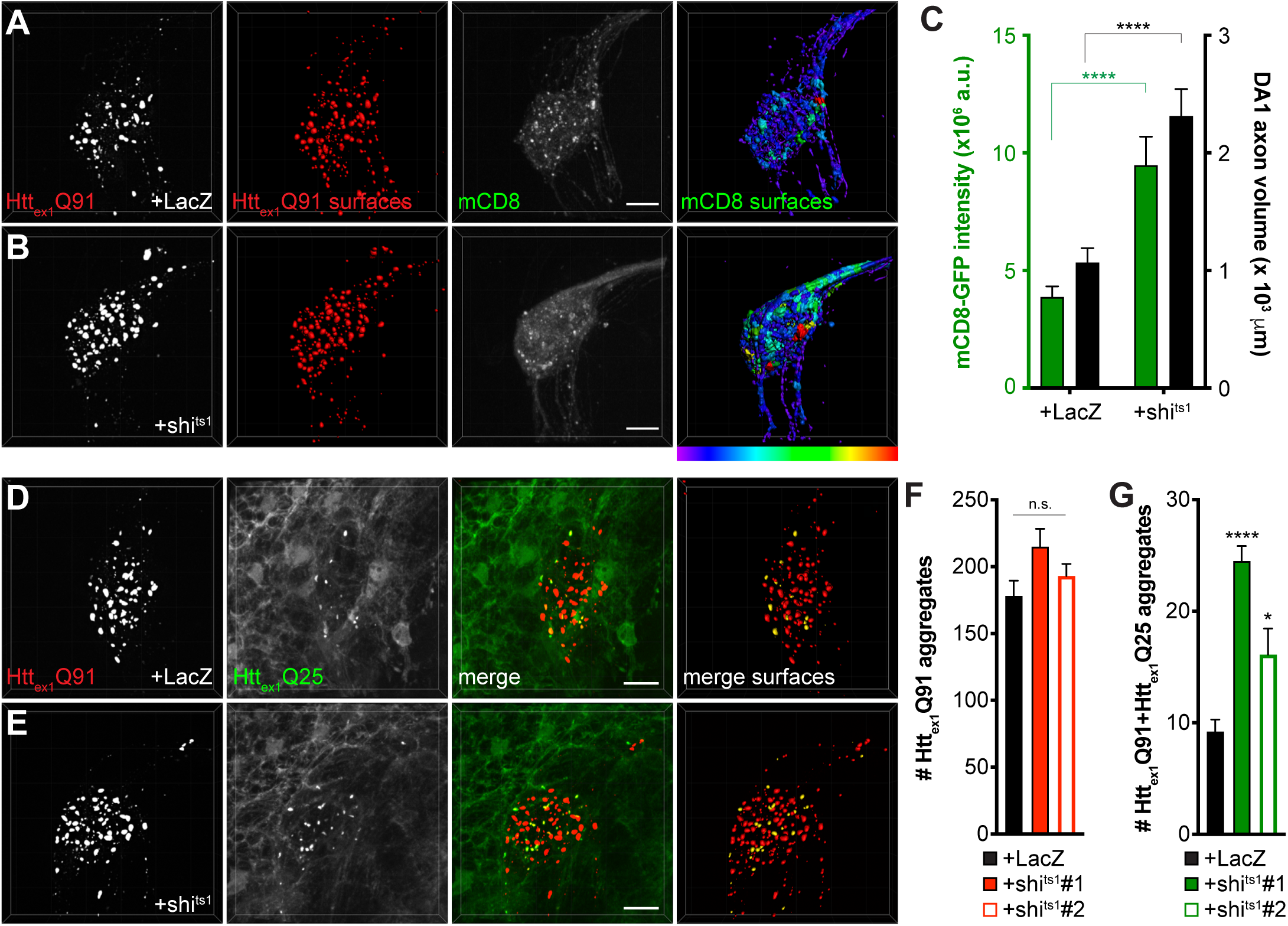
Inhibiting Shibire-mediated endocytosis increases mHttex1-expressing ORN axon volume and enhances transfer of mHtt_ex1_ aggregates from DA1 ORN axons to glia. (A and B) Confocal z-stacks of DA1 glomeruli from 10 day-old females co-expressing Htt_ex1_Q91-mCherry, mCD8- GFP, and either LacZ (A) or shi^ts1^ (B) in DA1 ORNs. Flies were shifted from the permissive temperature (18°C) to the restrictive temperature (31°C) upon eclosion. Raw data are shown in grayscale, and 3D segmented surfaces are shown in red for Htt_ex1_Q91 and as a heat map for mCD8-GFP to highlight differences in intensity between the genotypes. Scale bars = 10 μm. (C) Quantification of mCD8-GFP intensity (left y-axis, *green*) and volume (right y-axis, *black*) of DA1 glomeruli from 10 day-old adult females co-expressing LacZ or shi^ts1^ with Htt_ex1_Q91-mCherry and mCD8-GFP in DA1 ORNs. a.u. = arbitrary units. Data are shown as mean ± SEM; ****p < 0.0001 by Student’s t-test. (D and E) Confocal z-stacks of DA1 glomeruli from 5-6 day-old males expressing Htt_ex1_Q91-mCherry with either LacZ (D) or shi^ts1^ (E) in DA1 ORNs and Htt_ex1_Q25-YFP in repo+ glia. Adult flies were shifted from 18°C to 31°C upon eclosion. mCherry+ surfaces identified by semi-automated image segmentation are shown in the last panels, with Htt_ex1_Q91-only surfaces in red and Htt_ex1_Q91+Htt_ex1_Q25 surfaces in yellow. Scale bars = 10 μm. (F and G) Quantification of Htt_ex1_Q91 (F) and Htt_ex1_Q91+Htt_ex1_Q25 (G) aggregates in DA1 glomeruli of 5-6 day-old males expressing LacZ or shi^ts1^ using two independent *QUAS-shi^ts1^* lines. Data are shown as mean ± SEM; *p < 0.05, ****p < 0.0001 by one-way ANOVA with Tukey’s multiple comparisons test comparing shi^ts1^-expressing flies to control flies expressing LacZ.

To test whether inhibition of Shibire-mediated endocytosis in ORNs has similar effects on ORN-to-glia transfer of Htt_ex1_Q91 aggregates (Pearce et al., 2015), we co-expressed shi^ts1^ with Htt_ex1_Q91- mCherry in DA1 ORNs and monitored formation of seeded Htt_ex1_Q25 aggregates in the glial cytoplasm. Similar to effects of shi^ts1^ on trans-synaptic Htt_ex1_Q91 aggregate transfer, blocking Shibire-mediate endocytosis in DA1 ORNs increased seeded Htt_ex1_Q25 aggregate formation in glia without affecting Htt_ex1_Q91 aggregate numbers (Fig. 4D-G). Together, these findings suggest that shi^ts^-mediated silencing of Htt_ex1_Q91-mCherry-expressing DA1 ORNs alters axonal surface area and enhances prion-like transfer of Htt_ex1_Q91 aggregates from presynaptic ORN axons to the cytoplasm of both glia and postsynaptic PNs.

### Glial Draper is required for ORN-to-PN transfer and alters morphology of neuronal mHtt_ex1_ aggregates

The parallel effects of shi^ts1^-mediated endocytic blockade on transfer of Htt_ex1_Q91 aggregates from DA1 ORNs to DA1 PNs and to glia suggest that mHtt_ex1_ aggregate spreading between these different cell types is coordinated. Our prior work showed that ORN-to-glia transfer of mHtt_ex1_ aggregates is strictly dependent on Draper (Pearce et al., 2015), a scavenger receptor responsible for phagocytic engulfment and clearance of neuronal debris in the fly CNS and other tissues (Etchegaray et al.; Han et al., 2014; Hoopfer et al., 2006; MacDonald et al., 2006). Therefore, we sought to determine whether Draper-expressing phagocytic glia might play a role in transferring mHtt_ex1_ aggregates from ORNs to PNs. To test this, we quantified numbers of Htt_ex1_Q91 and seeded Htt_ex1_Q25 aggregates in DA1 ORN axons and PN dendrites, respectively, in animals heterozygous or homozygous for the *draper(drpr)^Δ5^* null mutation (Freeman et al., 2003). We previously reported that *drpr* knockout (KO) increased steady-state numbers of Htt_ex1_Q91 aggregates in DA1 ORN axons (Pearce et al., 2015); however, this effect was found to not be statistically significant between *drpr^Δ5^* heterozygotes and homozygotes in this study (Fig. 5A-C). We suspect this is the case for two reasons: (a) our image segmentation parameters improve identification of very small aggregates, which we show are more abundant when *drpr* is expressed at normal levels (Fig. 5E-G), and (b) the Htt_ex1_Q91-mCherry transgene used here expresses at lower levels than the transgene used in our previous study. Thus, Htt_ex1_Q91 aggregates initially form more slowly in DA1 ORN axons (compare numbers of Htt_ex1_Q91 aggregates in young females in Fig. 1J vs. Fig. 7I; the latter experiment used the same higher-expressing Htt_ex1_Q91-mCherry transgene as in our prior study) and may be less affected by Draper depletion. Strikingly though, *drpr* KO completely blocked formation of Htt_ex1_Q91+Htt_ex1_Q25 aggregates in postsynaptic PNs (Fig. 5A, B, and D), suggesting that Draper mediates trans-synaptic transfer of mHtt_ex1_ aggregates. This surprising effect of *drpr* KO on seeded Htt_ex1_Q25 aggregate formation was also observed when shi^ts1^ was co-expressed with Htt_ex1_Q91-mCherry in ORNs (Fig. 5-figure supplement 1), confirming that enhanced transfer of Htt_ex1_Q91 aggregates from shi^ts1^-expressing ORNs occurs via the same Draper-dependent mechanism. To rule out the possibility that *GH146-Gal4* drives expression of Htt_ex1_Q25-GFP in Draper+ glia, we used repo-Gal80 to inhibit Gal4 in all glia and saw no effect on Htt_ex1_Q91 or Htt_ex1_Q91+Htt_ex1_Q25 aggregate numbers (Fig. 5-figure supplement 2A, B, E, and F). Moreover, expression of *drpr*-specific siRNAs in PNs did not affect formation of either Htt_ex1_Q91 or Htt_ex1_Q91+Htt_ex1_Q25 aggregates (Fig. 5-figure supplement 2C-F), consistent with glia as the sole source of *drpr* expression in the fly CNS. We also did not observe significant colocalization between ORN-derived Htt_ex1_Q91 aggregates and GFP-fusions of Atg8a (Juhász et al., 2008) or Lamp1 (Pulipparacharuvil et al., 2005) in glia (Fig. 5-figure supplement 3), suggesting that neuronal Htt_ex1_Q91 aggregates do not seed Htt_ex1_Q25 in glial lysosomes or autophagosomes.

**Figure 5.**
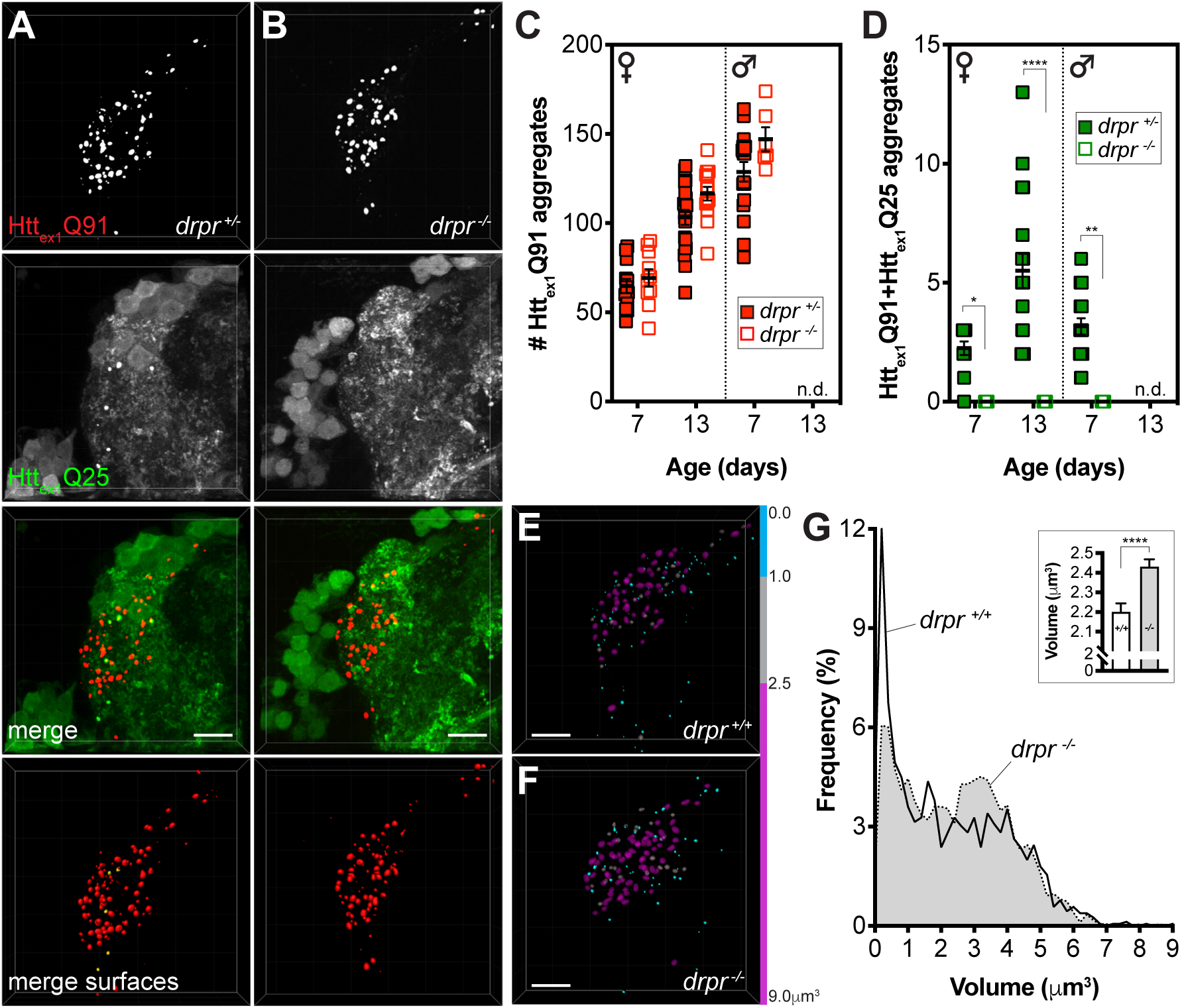
Draper mediates mHtt_ex1_ aggregate transfer from presynaptic DA1 ORNs to postsynaptic PNs and regulates neuronal mHtt_ex1_ aggregate size. (A and B) Confocal z-stacks of DA1 glomeruli from 13 day-old adult females expressing Htt_ex1_Q91-mCherry in DA1 ORNs and Htt_ex1_Q25-GFP in GH146+ PNs, either heterozygous (A; *drpr ^+/-^*) or homozygous (B; *drpr ^-/-^*) for the *drpr^Δ5^* null allele. mCherry+ surfaces identified by semi-automated image segmentation are shown in the last row, with Htt_ex1_Q91-only surfaces in red and Htt_ex1_Q91+Htt_ex1_Q25 surfaces in yellow. Scale bars = 10 μm. (C and D) Quantification of Htt_ex1_Q91 (C) and Htt_ex1_Q91+Htt_ex1_Q25 (D) aggregates in DA1 glomeruli from female or male *drpr ^+/-^* or *drpr ^-/-^* flies at the indicated ages. Data are shown as mean ± SEM; *p < 0.05, **p < 0.01, ****p < 0.0001 by two-way ANOVA with Sidak’s multiple comparisons test for *drpr ^+/-^* vs. *drpr ^-/-^* flies at the same ages. “n.d.” = not determined; 13 day-old *drpr ^-/-^* flies were not viable. (E and F) Htt_ex1_Q91 surfaces identified in DA1 glomeruli from 7 day-old *drpr ^+/-^* or *drpr ^-/-^* females expressing Htt_ex1_Q91-mCherry in DA1 ORNs. mCherry+ surfaces are color-coded according to the following volume ranges: cyan = 0-1.0 μm^3^; gray = 1.01-2.49 μm^3^; magenta = 2.5-9.0 μm^3^. Gray and magenta surfaces were set to 70% transparency to improve visibility of smaller cyan surfaces. Scale bars = 10 μm. (G) Relative frequency of volumes for all Htt_ex1_Q91 aggregates identified in 7 day-old *drpr ^+/+^* (*solid line*) or *drpr ^-/-^* (*dotted line; gray shading*) males and females. The inset graph shows mean Htt_ex1_Q91 aggregate volume ± SEM for the two genotypes. ****p < 0.0001 by unpaired Student’s t-test.

These results point to an unexpected but central role for glial Draper in Htt_ex1_Q91 aggregate transfer between multiple cell types in the fly CNS, but how phagocytic glia could mediate spreading of Htt_ex1_Q91 aggregates across neuronal synapses was not immediately clear. Intriguingly, mean volume of Htt_ex1_Q91 aggregates in DA1 ORN axons was significantly increased in *drpr* KO animals compared to wild-type controls (Fig. 5E, F, and G-inset), and the relative frequency of two aggregate subpopulations shifted between these genotypes: in the absence of *drpr*, the abundance of a smaller-sized subpopulation (∼0.1-1 μm^3^) decreased while a larger-sized subpopulation (∼2.5-4.0 μm^3^) increased in abundance (Fig. 5E-G). Intriguingly, the smaller aggregate subpopulation correlated well with the size of Htt_ex1_Q91 aggregates associated with converted Htt_ex1_Q25 in PNs (Fig. 2G) or glia (Pearce et al., 2015), suggesting that phagocytic glia could at least in part mediate formation of smaller seeding-competent Htt_ex1_ aggregates. However, molecular features other than size must regulate the seeding capacity of mHtt_ex1_ aggregates, since smaller-sized aggregates did not completely disappear in *drpr* KO animals (Fig. 5G). Taken together, these results indicate that Draper-expressing phagocytic glia mediate transfer of Htt_ex1_Q91 aggregates across DA1 ORN-PN synapses, perhaps in part by altering morphological features of neuronal mHtt_ex1_ aggregates.

### Caspase activation in ORNs is required for trans-synaptic transfer of mHtt_ex1_ aggregates

Phagocytic glia engulf injured or degenerating neuronal processes and apoptotic cell corpses by recognizing “eat me” signals exposed on these debris (Wilton et al., 2019), and mHtt expression induces caspase-dependent apoptosis in fly and mammalian models of HD (Ahmed et al., 2014). Thus, we asked whether Htt_ex1_Q91-mCherry-expressing DA1 ORNs activate pathways that could stimulate Draper-dependent transfer of aggregates to DA1 PNs. In *drpr^Δ5^* heterozygotes, expression of Htt_ex1_Q91- GFP in all ORNs did not significantly increase cleavage of *Drosophila* caspase-1 (Dcp-1) compared to flies expressing Htt_ex1_Q25-GFP (Fig. 6A, B, and E); however, Dcp-1 cleavage was significantly increased in Htt_ex1_Q91- vs. Htt_ex1_Q25-expressing ORNs when *drpr* was knocked out (Fig. 6C-E). These data suggest that glia efficiently clear Htt_ex1_Q91-expressing ORN axons displaying Dcp-1-dependent “eat me” signals via Draper-dependent phagocytosis. In addition, co-expression of the viral effector caspase inhibitor p35 (Hay et al., 1994) with Htt_ex1_Q91-mCherry in DA1 ORNs inhibited formation of Htt_ex1_Q91+Htt_ex1_Q25 aggregates in DA1 PNs (Fig. 6F, G, J, and K, *squares*), indicating that inhibition of apoptotic caspases in presynaptic ORNs phenocopies effects of *drpr* KO on Htt_ex1_Q91 aggregate transfer from ORNs to PNs. By contrast, co-expression of p35 with Htt_ex1_Q25-GFP in PNs did not affect numbers of Htt_ex1_Q91+Htt_ex1_Q25 aggregates in the DA1 glomerulus (Fig. 6H-K, *circles*), suggesting that caspase-dependent signaling in PNs are not required for Htt_ex1_Q91 aggregate spreading across ORN- PN synapses. Together, these results suggest that signals mediated by apoptotic caspase activation in DA1 ORNs, and not DA1 PNs, promote engulfment and trans-synaptic transfer of Htt_ex1_Q91 aggregates via phagocytic glia.

**Figure 6.**
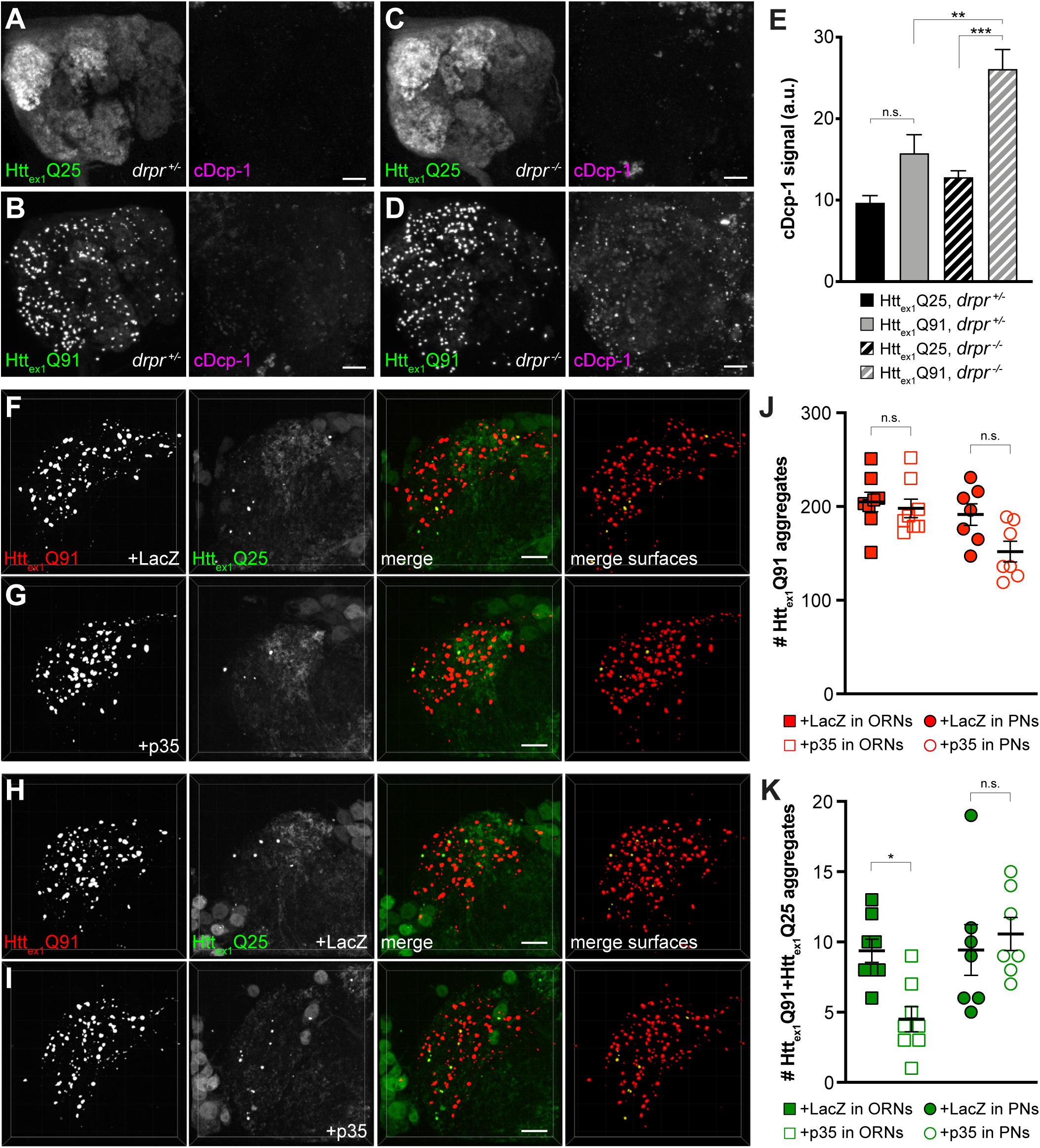
Caspase activation in ORNs mediates mHtt_ex1_ aggregate transfer from ORNs to PNs. (A-D) Maximum-intensity projections of antennal lobes from 8 day-old adult males expressing Htt_ex1_Q25-GFP (A and C) or Htt_ex1_Q91-GFP (B and D) in most ORNs using *Or83b-Gal4* in *drpr^Δ5^* heterozygotes (*drpr ^+/-^*; A and B) or homozygotes (*drpr ^-/-^*; C and D). Brains were immunostained for GFP (left panels) or cleaved Dcp-1 (right panels). Scale bars = 20 μm. (E) Quantification of cDcp-1 immunofluorescence from 8 day-old adult males with the same genotypes as in (A-D). Data are shown as mean ± SEM; **p < 0.01, ***p < 0.001, n.s. = not significant by one-way ANOVA with Tukey’s multiple comparisons test. (F-I) Confocal z-stacks of DA1 glomeruli from 14 day-old males expressing Htt_ex1_Q91-mCherry with LacZ (F) or p35 (G) in DA1 ORNs and Htt_ex1_Q25-GFP in GH146+ PNs, or Htt_ex1_Q91-mCherry in DA1 ORNs and Htt_ex1_Q25-GFP with LacZ (H) or p35 (I) in GH146+ PNs. mCherry+ surfaces identified by 3D segmentation are shown in the last panels, with Htt_ex1_Q91-only surfaces in red and Htt_ex1_Q91+Htt_ex1_Q25 surfaces in yellow. Scale bars = 10 μm. (J and K) Quantification of Htt_ex1_Q91 (J) or Htt_ex1_Q91+Htt_ex1_Q25 (K) aggregates in DA1 glomeruli of flies with the same genotypes in (F and G) (*squares*) or (H and I) (*circles*). Numbers of aggregates in flies expressing LacZ or p35 are indicated by solid or open shapes, respectively. Data are shown as mean ± SEM; *p < 0.05, n.s. = not significant by one-way ANOVA with Tukey’s multiple comparisons test.

### mHtt_ex1_ aggregates transfer from ORNs to PNs via the glial cytoplasm

The strict requirement for glial Draper in prion-like transfer of Htt_ex1_Q91 aggregates across ORN- PN synapses can be explained by two non-mutually exclusive models. In one model (Fig. 7A, *route 1*), mHtt_ex1_ aggregates spread from presynaptic ORNs to postsynaptic PNs via a glial cytoplasmic intermediate. This model is consistent with our previous finding that phagocytosed neuronal mHtt_ex1_ aggregates gain entry to the glial cytoplasm to effect prion-like conversion of Htt_ex1_Q25 (Pearce et al., 2015). Alternatively (Fig. 7A, *route 2*), phagocytic glia could sculpt the synaptic environment in a way that promotes transfer of mHtt_ex1_ aggregates directly from ORN axons to PN dendrites. To distinguish between these models, we generated transgenic flies that use three binary expression systems (i.e., QF-QUAS, Gal4-UAS, and LexA-LexAop) to independently express a uniquely-tagged Htt_ex1_ transgene in each of three cell populations: Htt_ex1_Q91-mCherry in DA1 ORNs, Htt_ex1_Q25-3xHA in repo+ glia, and Htt_ex1_Q25-YFP in GH146+ PNs (Fig. 7A). If Htt_ex1_Q91-mCherry aggregates formed in presynaptic ORNs transfer to postsynaptic PNs via the glial cytoplasm, Htt_ex1_Q91-mCherry aggregate seeds should template the aggregation first of Htt_ex1_Q25-3xHA in glia and then Htt_ex1_Q25-YFP in PNs, resulting in appearance of triple-labeled mCherry+/3xHA+/YFP+ puncta in the DA1 glomerulus (Fig. 7A, *route 1*). A transient double-labeled mCherry+/3xHA+ aggregate subpopulation that have accessed the glial cytoplasm but not yet reached PNs might also be observed in this scenario. Alternatively, if Htt_ex1_Q91- mCherry aggregates do not access the glial cytoplasm *en route* to PNs, only double-labeled mCherry+/3xHA+ and mCherry+/YFP+ aggregates would appear in the DA1 glomerulus (Fig. 7A, *route 2*).

**Figure 7.**
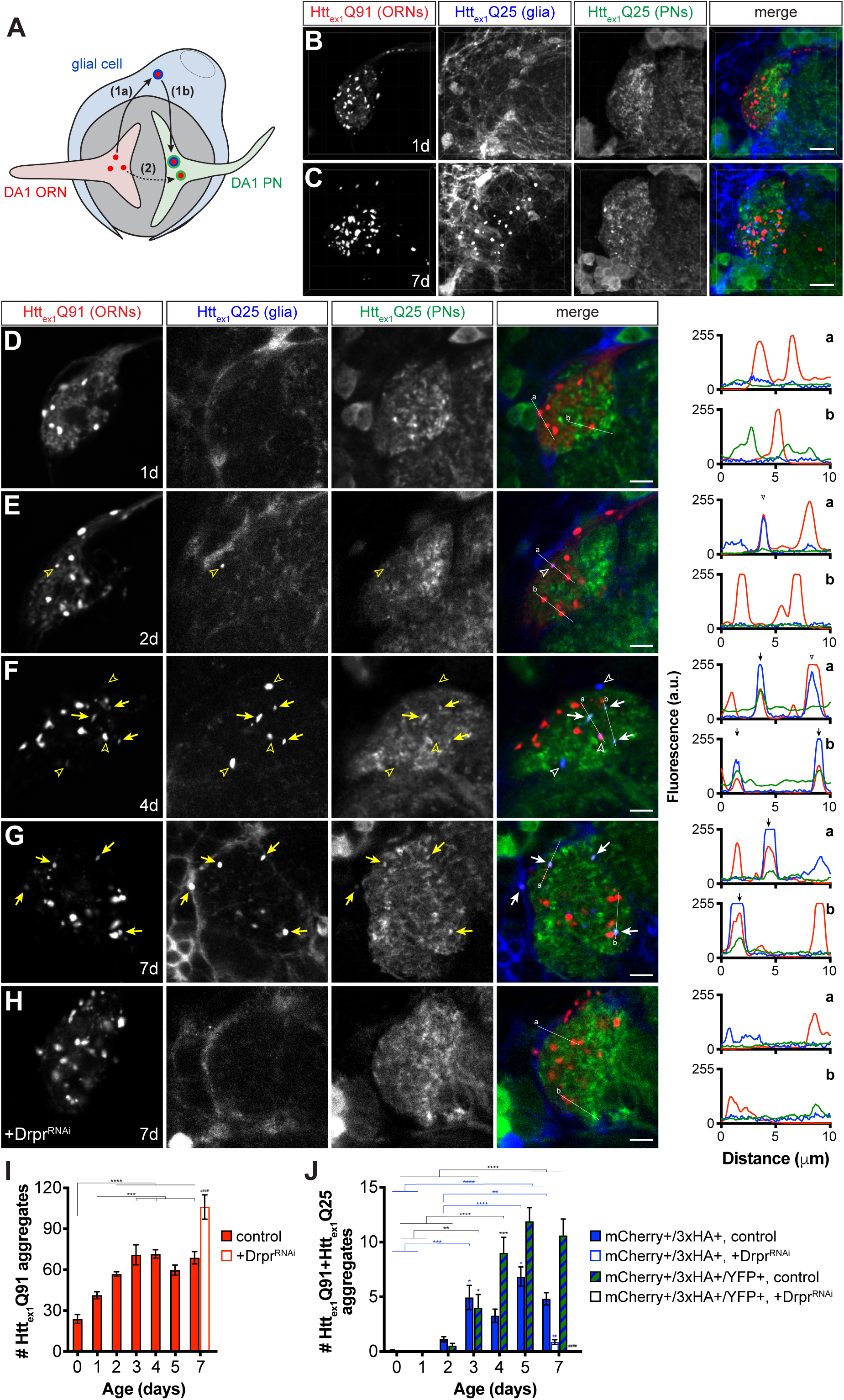
mHtt_ex1_ aggregates transfer from presynaptic ORNs to postsynaptic PNs via the cytoplasm of phagocytic glia. (A) Diagram illustrating our experimental approach for examining a role for Draper-expressing glia in mHtt_ex1_ aggregate transfer from DA1 ORNs to DA1 PNs. Flies that combined *Or67d-QF*-, *repo-Gal4*-, and *GH146-LexA:GAD*-driven expression of Htt_ex1_Q91-mCherry in DA1 ORNs, Htt_ex1_Q25-3xHA in all glia, and Htt_ex1_Q25-YFP in ∼60% of PNs, respectively, were generated and analyzed by confocal microscopy of immunostained brains. If Htt_ex1_Q91-mCherry aggregates travel to PNs via the glial cytoplasm (route 1), triple-labeled (mCherry+/3xHA+/YFP+) aggregates should be observed. By contrast, if Htt_ex1_Q91-mCherry aggregates transfer directly to PNs without accessing the glial cytoplasm (route 2), then only double-labeled (mCherry+/3xHA+ and mCherry+/YFP+) aggregates would be detected. (B-C) Confocal z-stacks of DA1 glomeruli from adult females of the indicated ages expressing Htt_ex1_Q91-mCherry in DA1 ORNs (red), Htt_ex1_Q25-3xHA in glia (blue), and Htt_ex1_Q25-YFP in PNs (green) at the indicated ages. Brains were immunostained with antibodies against the mCherry, 3xHA, and YFP tags unique to each Htt protein. Scale bars = 10 μm. (D-H) Single 0.35 μm confocal z-slices from females of the indicated ages with (D-G) the same genotype as in (B and C) or (H) also expressing dsRNAs targeting *drpr* (+Drpr^RNAi^). Colocalizing mCherry+/3xHA+ or mCherry+/3xHA+/YFP+ aggregates are indicated by open arrowheads or arrows, respectively, shown in yellow on grayscale and white on merged images for increased visibility. Scale bars = 5 μm. Htt_ex1_Q91-mCherry (red), Htt_ex1_Q25-3xHA (blue), and Htt_ex1_Q25-YFP (green) fluorescence intensity profiles for lines “a” and “b” are shown to the right of each merged image. Lines were scanned from leftmost to rightmost point. Arrowheads and arrows on graphs indicate peak mCherry fluorescence in colocalized mCherry+/3xHA+ and mCherry+/3xHA+/YFP+ aggregates, respectively. (I and J) Quantification of (I) mCherry-only or (J) mCherry+/3xHA+ and mCherry+/3xHA+/YFP+ aggregates identified in control (*solid bars*) or Drpr^RNAi^-expressing (*open bars*) animals over time. +Drpr^RNAi^ animals were only analyzed at 7 days-old. Data are shown as mean ± SEM; *p < 0.05, **p < 0.01, ***p < 0.001, or ****p < 0.0001 by one- or two-way ANOVA followed by Tukey’s multiple comparisons test. *s indicate statistical significance comparing control flies at different ages [black *s compare mCherry-only aggregates in (I) and mCherry+/3xHA+/YFP+ aggregates in (J), and blue *s compare mCherry+/3xHA+ aggregates in (J)], and #s indicate statistical significance comparing mCherry+/3xHA+ and mCherry+/3xHA+/YFP+ aggregates, respectively, in control vs Drpr^RNAi^-expressing flies at the same age.

Brains from transgenic flies expressing these differentially-tagged Htt_ex1_ transgenes in ORNs, glia, and PNs were analyzed, and expression patterns consistent with known morphologies of these cell types in the antennal lobe were observed by confocal microscopy (Fig. 7B and C, and Figure 7-figure supplement 1). In contrast to data in other figures, these samples required immunostaining to detect expression of Htt_ex1_Q25-3xHA and Htt_ex1_Q25-YFP expression in glia and PNs, respectively. We found that immunolabeled aggregate subtypes were not amenable to volumetric segmentation, and so we instead manually quantified single-, double-, and triple-labeled Htt_ex1_ aggregates in confocal slices and used line scan intensity profiling to confirm colocalization. Htt_ex1_Q91-mCherry expression in DA1 ORNs was partially diffuse and partially punctate in young (0-1 day-old) adult flies (Fig. 7B, D, E, and I), but became more punctate as the flies aged (Fig. 7C, F, G, and I). mCherry+/3xHA+ puncta representing Htt_ex1_Q91 aggregates that transferred from ORNs to glia first appeared in 2 day-old adults and increased in number over the next ∼24 hours (Fig. 7E, F, and J, *blue bars*). Remarkably, mCherry+/3xHA+/YFP+ puncta also began to appear in 2 day-old adults and increased in abundance until the flies were ∼4 days old (Fig. 7F, G, and J, *striped bars*). The timing of appearance of these different aggregate subtypes supports our hypothesis that Htt_ex1_Q91 seeds originating in DA1 ORN axons transit through the glial cytoplasm before reaching the cytoplasm of DA1 PN dendrites (Fig. 7A, *route 1*). mCherry+/YFP+ puncta representing aggregates that had transferred from DA1 ORNs to DA1 PNs without accessing the glial cytoplasm were not detected in these brains, arguing against direct transfer of Htt_ex1_Q91-mCherry aggregates from ORNs to PNs (Fig. 7A, *route 2*). Control animals expressing Htt_ex1_Q25 proteins in paired combinations of ORNs, glia, or PNs (Figure 7-figure supplement 1) confirmed that Htt_ex1_Q25 aggregates in glia or PNs were only detected when Htt_ex1_Q91 was expressed in ORNs. Remarkably, when we used RNAi to specifically knockdown *drpr* in glia, numbers of Htt_ex1_Q91-mCherry aggregates increased (Fig. 7H and I), while mCherry+/3xHA+ and mCherry+/3xHA+/YFP+ aggregates significantly decreased or were absent in 7 day-old adults (Fig. 7H and J). These data confirm that glial Draper mediates Htt_ex1_Q91 aggregate transmission from ORN axons to glia and across ORN-PN synapses. Taken together, these findings suggest that phagocytic glia act as obligatory intermediates in unidirectional prion-like spreading of mHtt_ex1_ aggregates from presynaptic ORNs to postsynaptic PNs in *Drosophila* brains.

## DISCUSSION

The hypothesis that prion-like spreading contributes to progression of protein aggregate pathology in HD and other neurodegenerative diseases is gaining considerable support, and yet we still understand very little about the mechanisms that underlie cell-to-cell spreading *in vivo*, the influences of cell- and tissue-specific vulnerability, and the relevance of aggregate transmission to disease progression. In this study, we report that mHtt_ex1_ aggregates transfer anterogradely from presynaptic ORNs to postsynaptic PNs via an obligatory path through phagocytic glia in *Drosophila* brains. ORN-to- PN transmission of mHtt_ex1_ aggregates was enhanced by blocking endocytosis and exocytosis and slowed by thermogenetically stimulating presynaptic ORNs, suggesting an inverse relationship between presynaptic activity and mHtt_ex1_ aggregate spreading. mHtt_ex1_ aggregate transfer across synapses was inhibited by blocking apoptosis in presynaptic neurons and required the Draper scavenger receptor, which we have previously reported to mediate phagocytic engulfment of mHtt_ex1_ aggregates from ORN axons and prion-like conversion of wtHtt_ex1_ in the glial cytoplasm (Pearce et al., 2015). Here, we expand our understanding of the role that glia play in prion-like diseases by showing that phagocytosed neuronal mHtt_ex1_ aggregates transit through the cytoplasm of glia before reaching postsynaptic PNs. To the best of our knowledge, these findings are the first to uncover a role for a well-conserved phagocytic pathway in prion-like spreading of pathogenic aggregates between neurons *in vivo*.

The increased propensity of mHtt to aggregate as a result of polyQ expansion is held in check by the proteostasis network, and intrinsic differences in proteostatic capacity of neuronal subpopulations could underlie regional selectivity to inclusion body formation in HD brains (Margulis and Finkbeiner, 2014). However, accumulating evidence that mHtt aggregates have prion-like properties suggests that aggregate spreading could also contribute to this by propagating mHtt pathology through networks of synaptically-connected but selectively-vulnerable neurons. We find that mHtt_ex1_ aggregate transmission across ORN-PN synapses occurs selectively in the anterograde direction and is inversely correlated with synaptic activity. Some of the earliest changes seen in HD patient brains involve loss of presynaptic cortical inputs to the striatum, where more prominent pathological findings (e.g., mHtt aggregate accumulation and massive loss of medium spiny neurons) appear in later stages of disease (Reiner and Deng, 2018). In addition, selective silencing of mHtt in both the cortex and striatum of BACHD mice inhibited striatal degeneration and motor phenotypes to a greater extent that silencing in just the striatum (Wang et al., 2014), suggesting that mHtt-induced toxicity in presynaptic cortical regions could play an important role in early HD development. Thus, presynaptic dysfunction may be a driving force for mHtt aggregate spreading between synaptically-connected regions of the brain. While our findings do not directly address secondary consequences of mHtt_ex1_ aggregate spreading across synapses, we identify a novel mechanism whereby glial responses to pathological changes in synapses mediate spreading of toxic aggregates between neurons. How aggregate spreading via glia impacts neuronal viability and functioning at the synaptic and/or circuit level are important questions for future studies.

Glia survey the brain to maintain homeostasis and can rapidly switch between supportive and reactive states in response to perturbations in CNS microenvironments. Upon sensing neuronal insult or injury, reactive microglia and astrocytes undergo dramatic morphological, metabolic, and transcriptional changes and promote neuronal survival by releasing trophic factors and phagocytosing debris (Hammond et al., 2018). Central roles for phagocytic glia in neurodegeneration are becoming increasingly recognized as genome-wide association studies and transcriptomic analyses identify glial genes associated with increased disease risk. For example, rare variants in the microglial phagocytic receptor gene *TREM2* are associated with increased risk of AD, FTD, and PD, and loss of TREM2 function exacerbates A*β*-, tau-, and *α*-synuclein-associated neurotoxicity (Griciuc et al.; Guo et al., 2019; Leyns et al., 2019; Zhao et al., 2018). Disruptions in key glial phagocytic functions leads to accumulation of potentially toxic aggregates in the brain (Asai et al., 2015; Hong et al., 2016; Pearce et al., 2015; Ray et al., 2017; Wilton et al., 2019), and administration of antibodies targeting pathological A*β*, tau, or *α*-synuclein proteins inhibits aggregate accumulation and spreading *in vivo* (Funk et al., 2015; Masliah et al., 2005; Tran et al., 2014; Yanamandra et al., 2013). However, chronically-active or otherwise dysfunctional glia induce neuroinflammation and exacerbate neuronal damage. For example, hyperactivate microglia excessively engulf synapses in pre-plaque AD mouse brains (Hong et al., 2016) and signal for formation of neurotoxic A1 astrocytes, which accumulate in several neurodegenerative diseases (Liddelow et al., 2017; Yun et al., 2018) and during aging (Clarke et al., 2018). Our findings thus add new insights to recognizing glia as double-edged players in neurodegeneration and suggest that rebalancing the protective and harmful effects of glial phagocytosis could be an effective new therapeutic strategy.

We provide several lines of evidence to support a model in which glia engulf mHtt_ex1_ aggregates, or perhaps portions of mHtt_ex1_ aggregate-containing axons, from ORNs without internalizing elements of PNs: (i) mHtt_ex1_ aggregate transmission occurred exclusively in the anterograde direction across ORN-PN synapses, (ii) neuronal mHtt_ex1_ aggregates did not colocalize with markers of lysosomes or autophagosomes in glia, and (iii) caspase inhibition in ORNs and not in PNs inhibited aggregate transfer. Our data suggest that glia selectively target presynaptic ORNs by recognizing apoptotic “eat me” signals induced by mHtt_ex1_ aggregate accumulation, and this process drives aggregate transfer to postsynaptic PNs. In developing mouse brains, pruned presynaptic structures are engulfed by microglia largely in the absence of postsynaptic markers (Schafer et al., 2012; Weinhard et al., 2018), suggesting that mammalian glia have the ability to selectively “nibble” presynaptic components (a fine-tuned phagocytic process known as trogocytosis). In addition, astrocytes internalize dystrophic presynaptic terminals that accumulate near amyloid plaques in transgenic mouse and human AD brains (Gomez-Arboledas et al., 2018). Intriguingly, astrocytes expressing the mammalian Draper homolog MEGF10 and complement-dependent microglia preferentially engulf “weaker” synapses to refine neural circuits in developing and adult mouse brains (Chung et al., 2013; Schafer et al., 2012), and aberrant activation of these pathways could contribute to early synaptic loss in neurodegenerative disease (Hong et al., 2016). We find that cell-to-cell mHtt_ex1_ aggregate transfer is accelerated from silenced ORN axons, suggesting that activity-impaired synapses are engulfed by nearby glia, enhancing spread along the ORN-to-glia-to-PN track. Thus, while the primary objective of phagocytic glia may be to eliminate toxic neuronal debris from the CNS, aberrant and/or excessive activation of phagocytic pathways could paradoxically promote disease. Premature elimination of live synapses could also drive network dysfunction, suggesting that attenuating glial responses could be an effective early intervention to preserve neurological functions in neurodegenerative disease patients (Carpanini et al., 2019).

The most exciting yet unexpected finding we report here is that prion-like mHtt_ex1_ aggregates do not directly transfer from ORNs to PNs, but instead make an obligatory detour through the phagocytic glial cytoplasm. This circuitous route could explain why relatively small numbers of seeded wtHtt_ex1_ aggregates form in glia or PNs, since seeding-competent mHtt_ex1_ must escape from multiple degradative systems and from the membrane-bound phagolysosomal compartment in order to access cytoplasmic wtHtt_ex1_. Though transfer through glia is compulsory across ORN-PN synapses, our data cannot exclude the possibility that mHtt_ex1_ aggregates transfer directly between neurons in other regions of the brain or at later stages of disease, when neuron-glia communication and cell integrity could be severely compromised. However, our findings suggest that phagocytic glia drive aggregate transfer across ORN-PN synapses by altering the transmissibility of mHtt_ex1_ aggregates formed in presynaptic ORN axons. We previously proposed that Draper-dependent phagocytosis could provide a temporary conduit for engulfed mHtt_ex1_ aggregates to escape into the glial cytoplasm (Pearce et al., 2015). This idea is supported by work from others showing that *α*-synuclein, tau, and mHtt_ex1_ aggregates rupture cell surface or endolysosomal membranes to access the cytoplasmic compartment (Chen et al., 2019; Falcon et al., 2018; Flavin et al., 2017; Ren et al., 2009; Zeineddine et al., 2015). It is possible that Draper-dependent phagocytosis could modify mHtt_ex1_ aggregates to increase their capacity to cross biological membranes, for example by altering molecular features such as rigidity, frangibility, or size. Indeed, we report here that seeding-competent mHtt_ex1_ aggregates belong to a smaller-sized aggregate subpopulation whose abundance is regulated by Draper, and other groups have observed that the seeding propensity of mHtt_ex1_ (Ast et al., 2018; Chen et al., 2001) or tau (Wu et al., 2013) aggregates is strongly associated with smaller size. This raises the intriguing possibility that membrane fission or fusion events that occur during dynamin-mediated engulfment or phagosome maturation could directly fragment or allow for escape of partially-digested neuronal mHtt_ex1_ aggregates that fall below an upper size limit for entry into the cytoplasm. Indeed, aggregate fragmentation and secondary nucleation events are thought to be key components of prion-like propagation in many neurodegenerative diseases (Knowles et al., 2014).

An outstanding question raised by our study is how ORN-derived mHtt_ex1_ aggregates physically transfer from glia to PN cytoplasms. This could be accomplished by a number of mechanisms already proposed for cell-to-cell spreading of prion-like aggregates, such as transport through extracellular vesicles or tunneling nanotubes (Costanzo et al., 2013; Sharma and Subramaniam, 2019), endocytosis/exocytosis (Asai et al., 2015; Babcock and Ganetzky, 2015; Chen et al., 2019; Holmes et al., 2013; Lee et al., 2010; Zeineddine et al., 2015), secretion or passive release of aggregates from dying cells, and direct penetration of lipid bilayers (Brundin et al., 2010; Davis et al., 2018; Vaquer-Alicea and Diamond, 2019). In a mouse model of tauopathy, pathological tau is transported between anatomically-connected regions of the brain via exosomes secreted by microglia (Asai et al., 2015), suggesting that phagocytosed neuronal aggregates may never encounter the extracellular space during transfer. It is also possible that dysfunction caused by continuous aggregate internalization, genetic mutations, and/or normal aging could decrease the efficiency by which glia clear aggregates, promoting their spread. In support of this, extracellular A*β* or *α*-synuclein fibrils accumulate inside microglia (Chung et al., 1999; Frackowiak et al., 1992), and the ability of microglia to effectively degrade phagocytosed material declines with age (Bliederhaeuser et al., 2016; Tremblay et al., 2012). We therefore favor a mechanism by which phagocytosed aggregates resistant to degradation overwhelm the glial phagolysosomal system and promote aggregate release, either through active secretion (e.g., in an act of self-preservation) or during glial cell death. While we did not observe mHtt_ex1_ aggregate transmission to non-partner PNs in flies up to 3 weeks old, suggesting that aggregate transfer at these ages is restricted to synaptic regions ensheathed by only 1-2 glial cells (MacDonald et al., 2006; Wu et al., 2017), important remaining questions are whether aggregates that have invaded the glial cytoplasm can transfer to other cells in or near the DA1 glomerulus (e.g., other glia or local interneurons) or to downstream neurons in the olfactory circuit.

In summary, our data demonstrate that phagocytic glia are active participants in the spread of prion-like protein aggregates between synaptically-connected neurons *in vivo*. Since microglial processes are highly motile (Hammond et al., 2018), these findings raise the intriguing possibility that phagocytic glia could mediate not only trans-synaptic, but also long-range spreading of neuronal aggregate pathology in the mammalian CNS. Our findings have important implications for understanding complex relationships between aggregate-induced cytotoxicity and neuron-glia communication in health and disease. Deciphering the mechanisms that regulate helpful vs. harmful effects of phagocytic glia in the brain will help to reveal the therapeutic potential of targeting key glial functions in HD and other neurodegenerative diseases.

## MATERIALS AND METHODS

### Fly husbandry

All fly stocks and crosses were raised on standard cornmeal/molasses media at 25°C, ∼50% relative humidity, and on a 12 hr light/12 hr dark cycle, unless otherwise noted. The following drivers were used to genetically access different cell populations in the fly CNS: *elav[C155]-Gal4* (Lin and Goodman, 1994), *Or67d-QF* (Liang et al., 2013), *GH146-Gal4* (Stocker et al., 1997), *GH146-QF* (Potter et al., 2010), and *GH146-LexA::GAD* (Lai et al., 2008) (a kind gift from Tzumin Lee, Janelia Farms), *repo-Gal4* (Sepp et al., 2001), and *Or83b-Gal4* (Kreher et al., 2005). Transgenic flies previously described by our lab include *QUAS-Htt_ex1_Q91-mCherry*, *QUAS-Htt_ex1_Q25-mCherry*, *UAS-Htt_ex1_Q91-mCherry*, *UAS-Htt_ex1_Q25-GFP*, *UAS-Htt_ex1_Q25-YFP*, and *UAS-GFP* transgenes inserted at the attP3 (1^st^ chromosome) and/or attP24 (2^nd^ chromosome) ϕC31 integration sites (Pearce et al., 2015). Other transgenic flies not generated in this study include: *UAS-mCD8-GFP* (BDSC #5137), *QUAS-nucLacZ* lines #7 and 44 (BDSC #30006 and 30007)*, QUAS-shi^ts1^* lines #2, 5, and 7 (BDSC #30010 and 30012; kind gift from C. Potter, Johns Hopkins School of Medicine), *QUAS-TeTxLC* lines #4c and 9c and *QUAS-dTrpA* lines #5, 6, and 7 (kind gifts from O. Riabinina and C. Potter, Johns Hopkins), *QUAS-mCD8-GFP* line #5J (BDSC #30002), *QUAS-p35* (a kind gift from H. Steller, Rockefeller University), *UAS-LacZ* (BDSC #8529), *UAS-p35* (BDSC #5072)*, QUAS-Gal80* (BDSC #51948), *UAS-QS* (BDSC #30033), *repo-Gal80* (a kind gift from T. Clandinin, Stanford University), *UAS-FFLuc.VALIUM1* (BDSC #35789), *UAS-GFP-Lamp1* (BDSC #42714), *UAS-Atg8a-GFP* (BDSC #52005), and *UAS-Draper^RNAi^* and *drpr^Δ5^* mutant flies (kind gifts from M. Freeman, Vollum Institute). Genotypes for all flies used in this study are listed in Table 1.

**Table.**
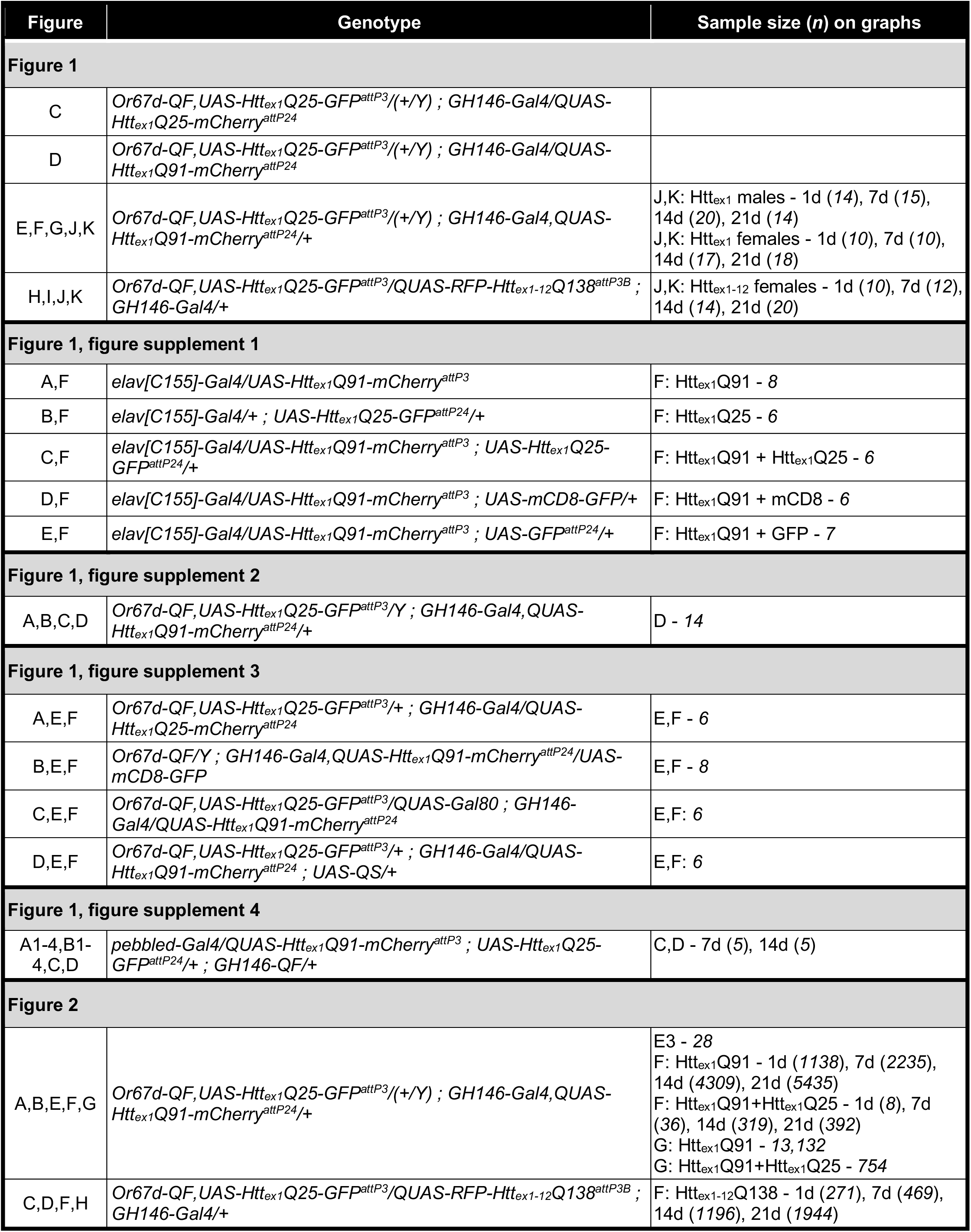
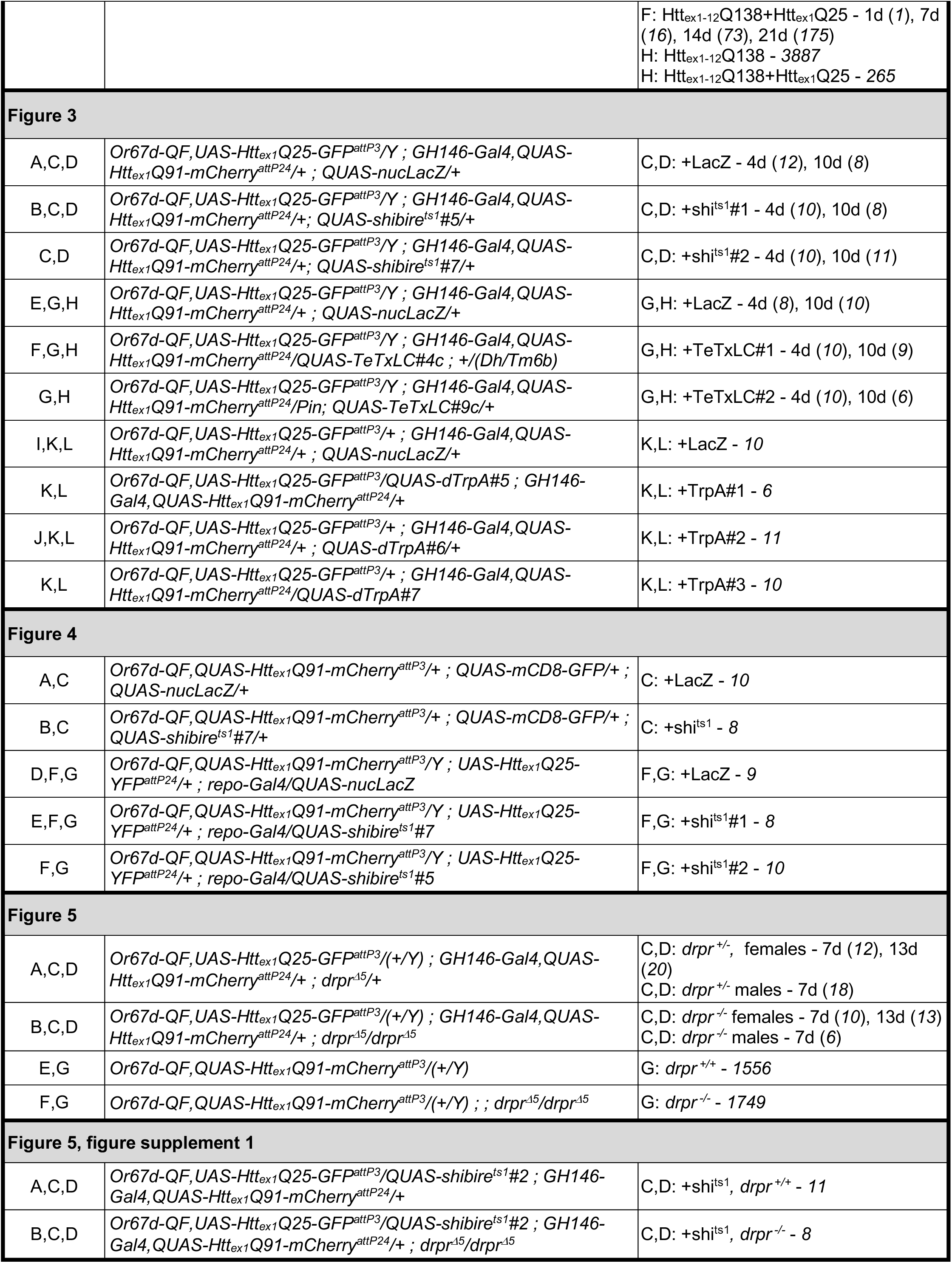
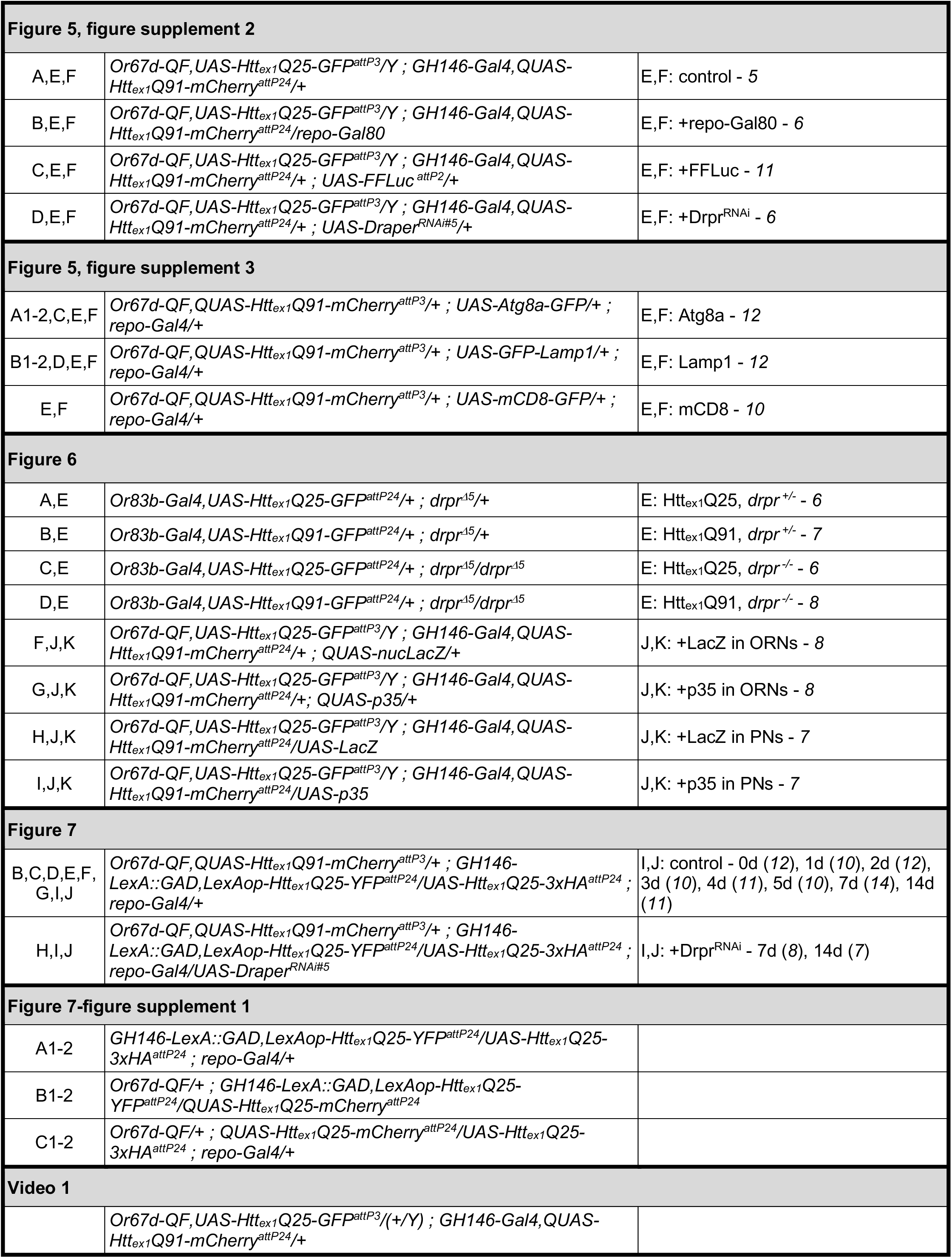
Genotypes and sample sizes displayed in each figure

### Cloning and transgenesis

pUASTattB(Htt_ex1_Q25-GFP) and pUASTattB(Htt_ex1_Q25-3xHA) plasmids were generated by PCR amplification of tagged Htt_ex1_ cDNAs from the pcDNA3 vector backbone and cloned into pUASTattB via XhoI and XbaI restriction sites. cDNA for mRFP-Htt_ex1-12_Q138 (a kind gift from T. Littleton, MIT) (Weiss et al., 2012) was subcloned into the pQUASTattB plasmid (Riabinina et al., 2015) using EcoRI and XbaI restriction sites. Htt_ex1_Q25-YFP cDNA was amplified by PCR from the pUASTattB(Htt_ex1_Q25-YFP) plasmid (Pearce et al., 2015) and subcloned into the EcoRI site in pLOT downstream of LexA-responsive LexAop DNA sequences. Plasmids were microinjected into embryos containing the attP3 X chromosome (for UAS-Htt_ex1_Q25-GFP), attP3B or attP8 X chromosome (for QUAS-mRFP-Htt_ex1- 12_Q138), or attP24 2^nd^ chromosome (for UAS-Htt_ex1_Q25-3xHA or LexAop-Htt_ex1_Q25-YFP) ϕC31 integration sites either in-house or at BestGene, Inc (Chino Hills, CA).

### Drosophila brain dissection and sample preparation

Adult fly brains were dissected, fixed, and stained and/or imaged as previously described (Pearce et al., 2015). Briefly, brains were dissected in ice-cold phosphate-buffered saline containing either 0.03% Triton X-100 (PBS/0.03T, when intrinsic fluorescence of FP-fusions was imaged) or 0.3% Triton X-100 (PBS/0.3T, when indirect immunofluorescence was used to detect protein expression). Where possible, we imaged using intrinsic GFP/YFP/mCherry fluorescence when Htt-FP fusions expression levels were high enough to detect on the confocal. Dissected brains were transferred to microfuge tubes containing PBS/T + 4% paraformaldehyde fixative solution on ice and then fixed in the dark at room temperature (RT) for 5 min (when imaging intrinsic fluorescence) or 20 min (when using immunofluorescence). For direct fluorescence imaging, brains were washed in PBS/0.03T buffer several times before incubation in Slowfade Gold Antifade Mountant (Invitrogen, Carlsbad, CA) for at least 1 hr at 4°C in the dark. For immunostaining, brains were washed several times in PBS/0.3T, and then blocked in PBS/0.3T containing 5% normal goat serum (Lampire Biological Laboratories, Pipersville, PA) for 30min at RT, followed by incubation in primary antibodies diluted in blocking solution and incubation for 24-72 hr at 4°C in the dark. Brains were then washed in PBS/0.3T several times at RT and incubated in secondary antibodies diluted in blocking solution for 20-24 hr at 4°C in the dark. Following another set of washes in PBS/0.3T at RT, the brains were incubated in Slowfade mountant for at least 16 hr at 4°C in the dark. Brains were then bridge-mounted in Slowfade mountant on glass microscopy slides overlayed with #1.5 coverglass (22 x 22 mm), and edges were sealed using clear nail polish.

Primary antibodies used in this study include rabbit anti-DsRed (1:2000; #632496; Takara Bio USA, Inc., Mountain View, CA), rabbit anti-mCherry (1:500; #PA5-34974; Invitrogen, Carlsbad, CA), chicken anti-GFP (1:500; #GFP-1020; Aves Labs, Tigard, OR), chicken anti-GFP (1:1000; #ab13970; Abcam, Cambridge, UK), chicken anti-GFP (1:500; #A10262; Invitrogen, Carlsbad, CA), rat anti-HA (1:100; clone 3F10; #11867423001; Roche, Basel, Switzerland), rabbit anti-cleaved Dcp-1 (1:100; #9578S; Cell Signaling Technology, Danvers, MA), and mouse anti-Bruchpilot (1:100; clone nc82; Developmental Studies Hybridoma Bank, Iowa City, IA). Secondary antibodies used include FITC- conjugated donkey anti-chicken (1:200; #703-095-155; Jackson Immuno Research Labs, West Grove, PA) and AlexaFluor 488, 568, or 647-conjugated goat anti-chicken, anti-mouse, anti-rabbit, or anti-rat IgGs (1:250 each; Invitrogen, Carlsbad, CA).

### Image acquisition

All data were collected on a Leica SP8 laser-scanning confocal system equipped with 405 nm, 488 nm, 561 nm, and 633 nm lasers and 40X 1.3NA or 63X 1.4NA oil objective lenses. Leica LAS X software was used to establish optimal settings during each microscopy session and to collect optical z-slices of whole-mounted brain samples with Nyquist sampling criteria. Optical zoom was used to further magnify and establish regions of interest in each sample. For most images, confocal data were collected from ∼60 x 60 x 25 μm (*xyz*) stacks centered on a single DA1 glomerulus, which was located using fluorescent signal from Htt_ex1_ protein expressed in DA1 ORN terminals. Exceptions to this are shown in Fig. 1-figure supplement 1A-E [250 x 250 x ∼60 μm (*xyz*) stacks capturing fluorescence signal in the anterior central brain] and Fig. 1C-D, Fig. 1-figure supplement 4A-B, and Fig. 6A-D [∼150 x 150 x 30 μm (*xyz*) stacks of the anterior portion of a single antennal lobe].

### Post-imaging analysis

Raw confocal data were analyzed in 2D using ImageJ/FIJI (NIH, Bethesda, MD) or in 3D using Imaris (Bitplane, Zürich, Switzerland), and all quantitative data were analyzed independently by two researchers blinded to the experimental conditions. For semi-automated quantification of aggregates in DA1 glomeruli, raw confocal data were deconvolved to reduce blur, rendered in 3D, and cropped if necessary to establish the region of interest for further analysis. mHtt fluorescence was segmented in 3D stacks using the “Surfaces” algorithm (surface detail set to 0.25 μm and background subtraction at 0.75 μm), with the “split touching objects” option selected, and seed point diameter was set to 0.85 μm. Background thresholding and seed point classification were adjusted manually for each image to optimize segmentation of heterogeneously-sized Htt_ex1_Q91-mCherry puncta (“aggregates”) and minimize capturing of diffuse signal. These settings differed <5-10% among individual samples in the same experiment. In rare cases (<5% of aggregates in each image), some larger aggregates were aberrantly split or smaller aggregates in close proximity were incorrectly merged; in these cases, the objects were manually unified or split using the software program. To quantify seeded wtHtt_ex1_ aggregates, red mHtt surfaces that colocalized with Htt_ex1_Q25-GFP were identified by applying a filter for mean intensity in the green channel. Colocalizing aggregates were selected by adjusting the threshold to capture discrete Htt_ex1_Q25 puncta with high contrast compared to surrounding diffuse signal. Segmentation of Htt_ex1_Q25 fluorescence in Fig. 1-figure supplement 2, Fig. 2A-D, and Video 1 was carried out using the same settings described above in the green channel. To measure DA1 ORN axon volume and intensity in Fig. 4A-C, mCD8-GFP fluorescence was segmented using the “Surfaces” function in Imaris, with surface detail set to 0.2 μm and background subtraction at 3 μm. Detailed surface measurements (e.g. volume or intensity) were calculated in Imaris, and the data were exported to Excel (Microsoft Corporation, Redmond, WA) or Prism (GraphPad Software, San Diego, CA) for further analysis.

For data shown in Fig. 7 and Fig. 7-figure supplement 1, indirect immunofluorescence was used to detect Htt_ex1_Q25-3xHA expression in glia and amplify low Htt_ex1_Q25-YFP signal in PNs. We found that semi-automated image segmentation as described above reported fewer than half of immunolabeled colocalized aggregates identified by manual counting, likely because of poor antibody penetration into the aggregate core and Htt_ex1_Q25-YFP aggregates with low contrast that made these data less amenable to 3D segmentation. Instead, we manually counted aggregates in these samples by scanning individual z-slices and scoring discrete puncta with increased signal relative to adjacent diffuse signal in one, two, or all three channels. Line scans confirmed colocalization of signals from the different Htt_ex1_ proteins, as shown in Fig. 7D-H.

### FRET analysis

Htt_ex1_Q25+Htt_ex1_Q91 colocalized aggregates were analyzed for FRET using the acceptor photobleaching method as previously described (Pearce et al., 2015). Briefly, 28 individual aggregates were analyzed by photobleaching the mCherry acceptor appended to Htt_ex1_Q91 using a 561nm laser set at 100% intensity and scanning until fluorescence was no longer detectable. Donor fluorescence dequenching was measured by exciting GFP fused to Htt_ex1_Q25 using a 488 nm laser set at 1% intensity before and after acceptor photobleaching. mCherry fluorescence was also excited before and after photobleaching with a 561 nm laser set at 1% intensity. Fluorescence emission was collected between 500 and 550 nm for GFP and 610 and 700 nm for mCherry to generate before and after images as shown in Fig. 2E. FRET efficiencies (FRET_eff_) were calculated after background correction using the equation (GFP_after_-GFP_before_)/(GFP_after_) x 100 and represented by a pixel-by-pixel FRET_eff_ image generated using the FRETcalc plugin (Stepensky, 2007) in FIJI/ImageJ.

### Statistical analyses

All quantified data were organized and analyzed in Excel or Prism 8. Quantifications in graphical form are shown as mean ± SEM, except for frequency analyses, which are displayed in histograms. Results of all statistical analyses are described in each Figure Legend. Sample size (*n*) for each figure is indicated in Table 1 and was selected to yield sufficient statistical power (*≥* 5 biological replicates from *≥* 3 brains for each condition; a single DA1 glomerulus represents one biological replicate). Multiple statistical comparisons were performed using the following tests and post-hoc corrections where appropriate: Student’s *t-*tests for pairwise comparisons, or one-way or two-way ANOVA followed by Sidak’s or Tukey’s multiple comparison tests for experiments involving *≥*3 genotypes. Results of these statistical tests are shown on each graph, and symbols used to indicate statistical significance are defined in the Figure Legends.

## ACKNOWLEDGMENTS

The authors would like to thank K. Chao, D. Luginbuhl, V. Reed, B. Temsamrit, and M. Warkala for technical assistance, G. Panning for help with analyzing detailed aggregate measurements, T.R. Clandinin, M.R. Freeman, H. Krämer, T. Lee, J.T. Littleton, C.J. Potter, and O. Riabinina for kindly sharing reagents, and members of the Kopito, Luo, and Pearce labs for many valuable discussions. This work was supported by grants from the NIH (R01-DC005982 to L.L., R01-NS042842 to R.R.K, and R03-AG063295 to M.M.P.P.), the W.W. Smith Charitable Trusts (to M.M.P.P.), the Pittsburgh Foundation (UN2018-98318 to M.M.P.P.), and start-up funds from University of the Sciences (to M.M.P.P.).

## AUTHOR CONTRIBUTIONS

K.M.D., O.R.D., A.D.A.Z., G.E.P., and W.M.T. – Investigation, Validation, Visualization, Writing – review & editing. L.L. and R.R.K. – Conceptualization, Resources, Supervision, Writing – review & editing, Funding Acquisition. M.M.P.P. – Conceptualization, Methodology, Investigation, Validation, Formal Analysis, Visualization, Writing – original draft preparation, Writing – review & editing, Resources, Supervision, Project Administration, Funding Acquisition.

## FIGURE LEGENDS

**Figure 1 – figure supplement 1.**
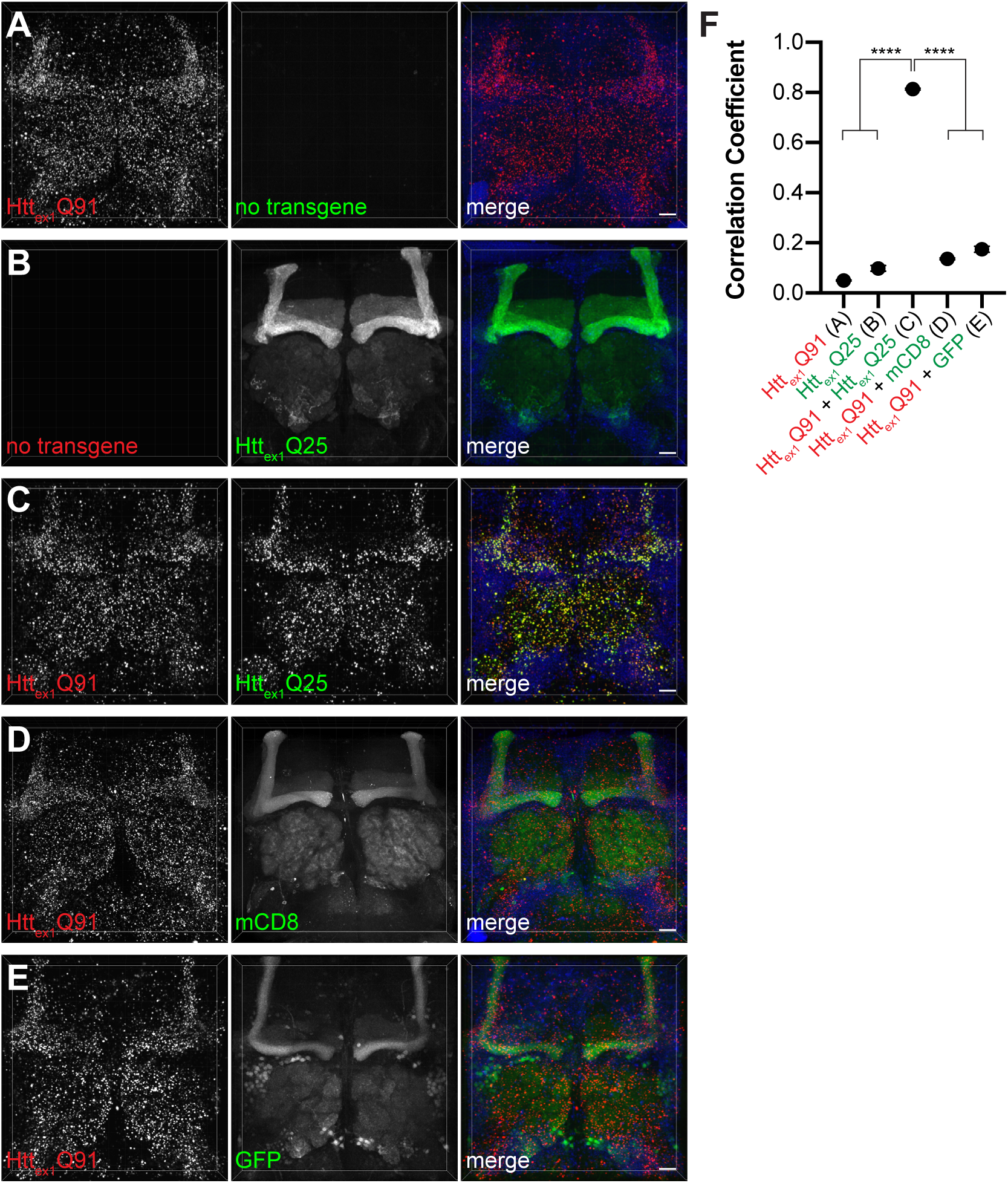
mHtt_ex1_ nucleates prion-like conversion of wtHtt_ex1_ in fly neurons. (A-E) Confocal z-stacks of 14 day-old adult female brains expressing (A) Htt_ex1_Q91-mCherry or (B) Htt_ex1_Q25-GFP alone, or co-expressing Htt_ex1_Q91-mCherry with (C) Htt_ex1_Q25-GFP, (D) membrane-targeted mCD8-GFP, or (E) soluble GFP in all neurons using *elav[C155]-Gal4*. Dimensions of each confocal stack are 250 x 250 x ∼60 (*xyz*) μm. Merged images include DAPI signal (*blue*) to label nuclei. Scale bars = 20 μm. (F) Colocalization of mCherry and GFP fluorescent signals calculated as Pearson’s correlation coefficients for genotypes shown in (A-E). Data are shown as mean ± SEM; ****p < 0.0001 by one-way ANOVA with Tukey’s multiple comparisons test.

**Figure 1 – figure supplement 2.**
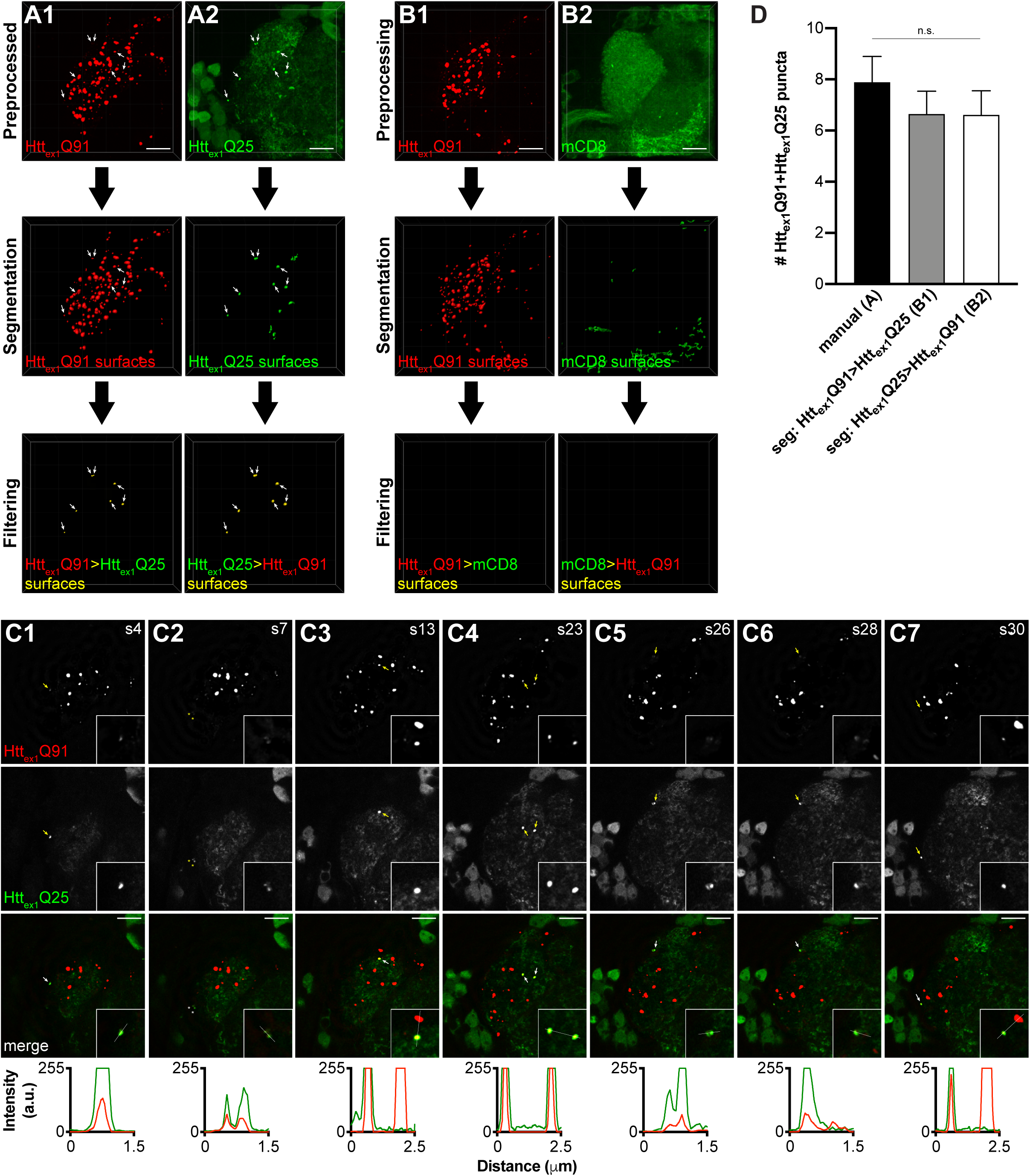
Semi-automatic quantification of seeded wtHtt_ex1_ aggregates. (A and B) Semi-automatic image processing workflow for high-magnification confocal z-stacks of DA1 glomeruli from 14 day-old adult males expressing Htt_ex1_Q91-mCherry in DA1 ORNs and either (A1,2) Htt_ex1_Q25-GFP or (B1,2) mCD8-GFP in GH146+ PNs. Raw data were preprocessed by deconvolution to reduce noise (*top panels*), segmented in the mCherry (A1 and B1) or GFP (A2 and B2) channels (*middle panels*), and filtered for co-localizing fluorescence signal in the other channel (*bottom panels*). Arrows in (A1 and A2) indicate seven Htt_ex1_Q91+Htt_ex1_Q25 puncta identified by this method. Scale bars = 10 μm. (C1-7) Selected single 0.35 μm confocal z-slices from the same confocal stack shown in (A1 and A2). Slice number is indicated at the top right of each image. Individual Htt_ex1_Q91+Htt_ex1_Q25 puncta identified by semi-automated image segmentation in (A) are indicated with arrows (*yellow* in individual channels, *white* on merged images) in each slice. Two additional co-localized Htt_ex1_Q91+Htt_ex1_Q25 puncta identified by manual counting are indicated with asterisks in (A2). Scale bars = 10 μm. Insets show Htt_ex1_Q91+Htt_ex1_Q25 puncta at higher zoom (inset dimensions = 9.12 μm x 9.12 μm). Htt_ex1_Q91-mCherry (*red*) and Htt_ex1_Q25-GFP (*green*) fluorescence intensity profiles for lines indicated in merged insets are shown below images. Lines were scanned from leftmost to rightmost point. (D) Comparison of manual quantification (C) vs semi-automated segmentation approaches (A1 and A2) for 14 day-old males with the same genotype in (A and C). Data are shown as mean ± SEM; n.s. = not significant by one-way ANOVA followed by Tukey’s multiple comparisons test.

**Figure 1 – figure supplement 3.**
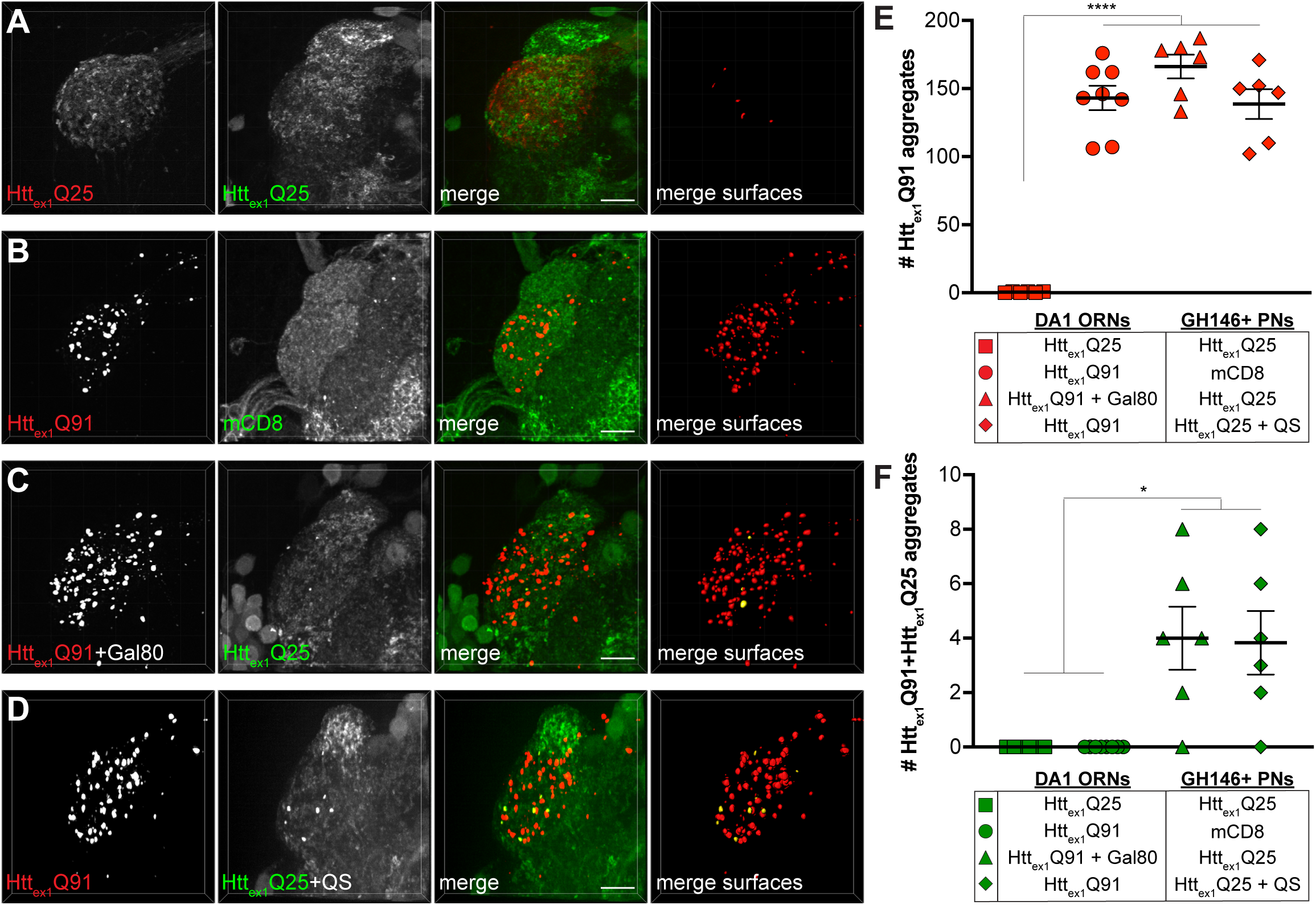
Controls for prion-like transmission of mHtt_ex1_ aggregates from presynaptic DA1 ORNs to postsynaptic PNs. (A-D) Confocal z-stacks of DA1 glomeruli from 10 day- old adults expressing (A) Htt_ex1_Q25-mCherry in DA1 ORNs and Htt_ex1_Q25-GFP in GH146+ PNs, (B) Htt_ex1_Q91-mCherry in DA1 ORNs and mCD8-GFP in GH146+ PNs, (C) Htt_ex1_Q91-mCherry together with Gal80 in DA1 ORNs and Htt_ex1_Q25-GFP in GH146+ PNs, and (D) Htt_ex1_Q91-mCherry in DA1 ORNs and Htt_ex1_Q25-GFP together with QS in GH146+ PNs. mCherry+ surfaces identified by semi-automated image segmentation are shown in the last panels, with Htt_ex1_Q91-only surfaces in red and Htt_ex1_Q91+Htt_ex1_Q25 surfaces in yellow. Scale bars = 10 μm. (E and F) Quantification of (E) Httex1Q91 and (F) Htt_ex1_Q91+Htt_ex1_Q25 aggregates in the DA1 glomeruli of flies with genotypes shown in (A-D). Data are shown as mean ± SEM. *p < 0.05, ****p < 0.0001 by one-way ANOVA with Tukey’s multiple comparisons test.

**Figure 1 – figure supplement 4.**
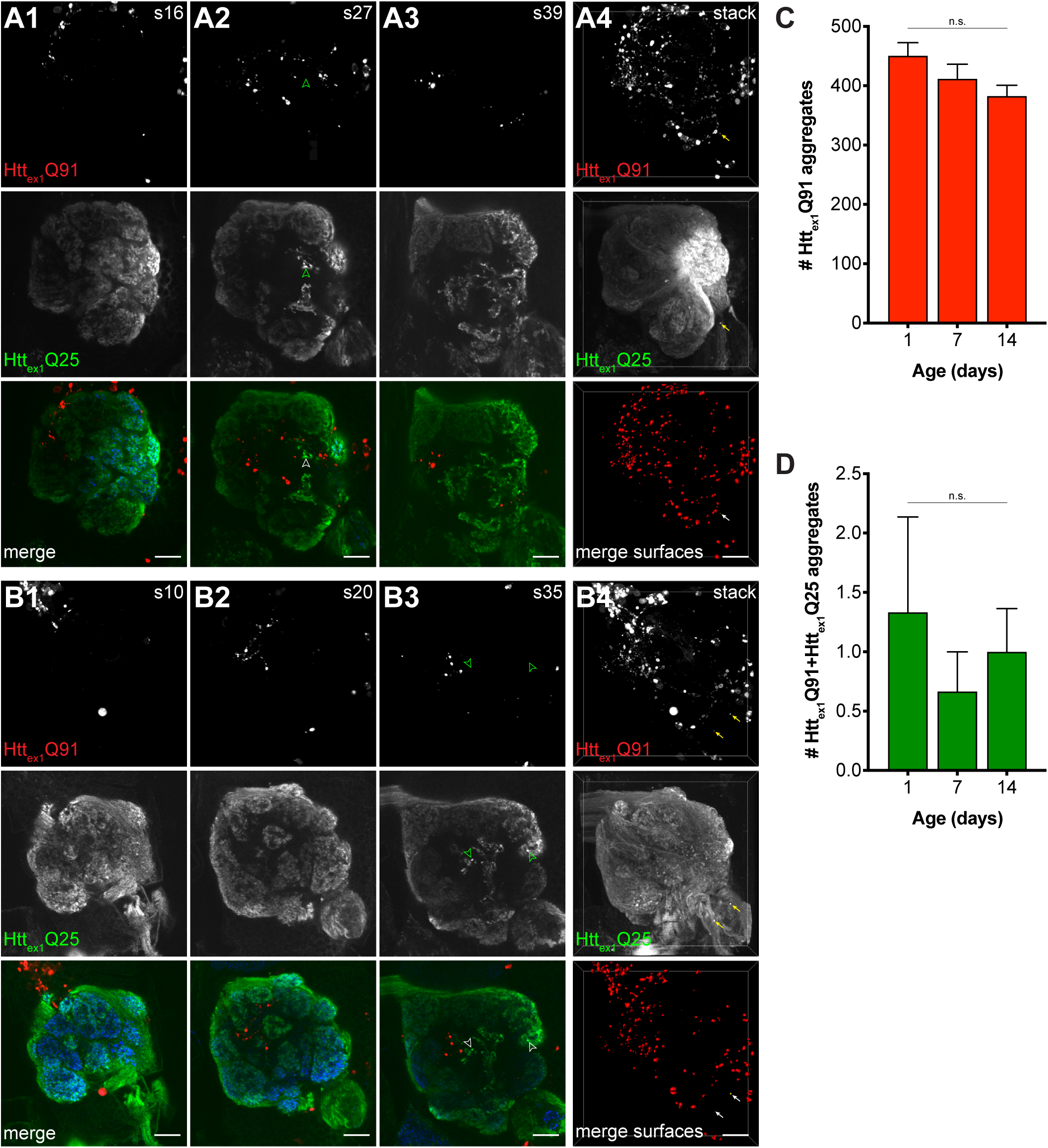
mHtt_ex1_ aggregates do not transfer retrogradely from PN dendrites to ORN axons. Single confocal slices (A1-3 and B1-3) and confocal z-stacks (A4 and B4) of the antennal lobe from 7 (A) and 14 (B) day-old adult females expressing Htt_ex1_Q91-mCherry in ∼60% of PNs using *GH146-QF* and Htt_ex1_Q25-GFP in all ORNs using *pebbled-Gal4*. Dissected brains were immunostained with antibodies against mCherry (*red*), GFP (*green*), and the neuropil marker Bruchpilot (*blue*, shown in merged images). GFP+ puncta identified in single slices are indicated by open arrowheads; none of these were found to be mCherry+. Semi-automated segmentation of the Htt_ex1_Q91-mCherry fluorescent signal (“merge surfaces” in A4 and B4) identified numerous Htt_ex1_Q91 aggregates (graphed in C) throughout the antennal lobe neuropil and surrounding region. A small number of Htt_ex1_Q91+Htt_ex1_Q25 surfaces (graphed in D) were identified in these brains (arrows in A4 and B4); however, none of these were located within the boundaries of the antennal lobe. Scale bars = 10 μm; slice numbers are indicated at the top right in (A1-3 and B1-3). Quantified data in (C and D) are shown as mean ± SEM; n.s. = not significant by one-way ANOVA with Tukey’s multiple comparisons test.

**Figure 5 – figure supplement 1.**
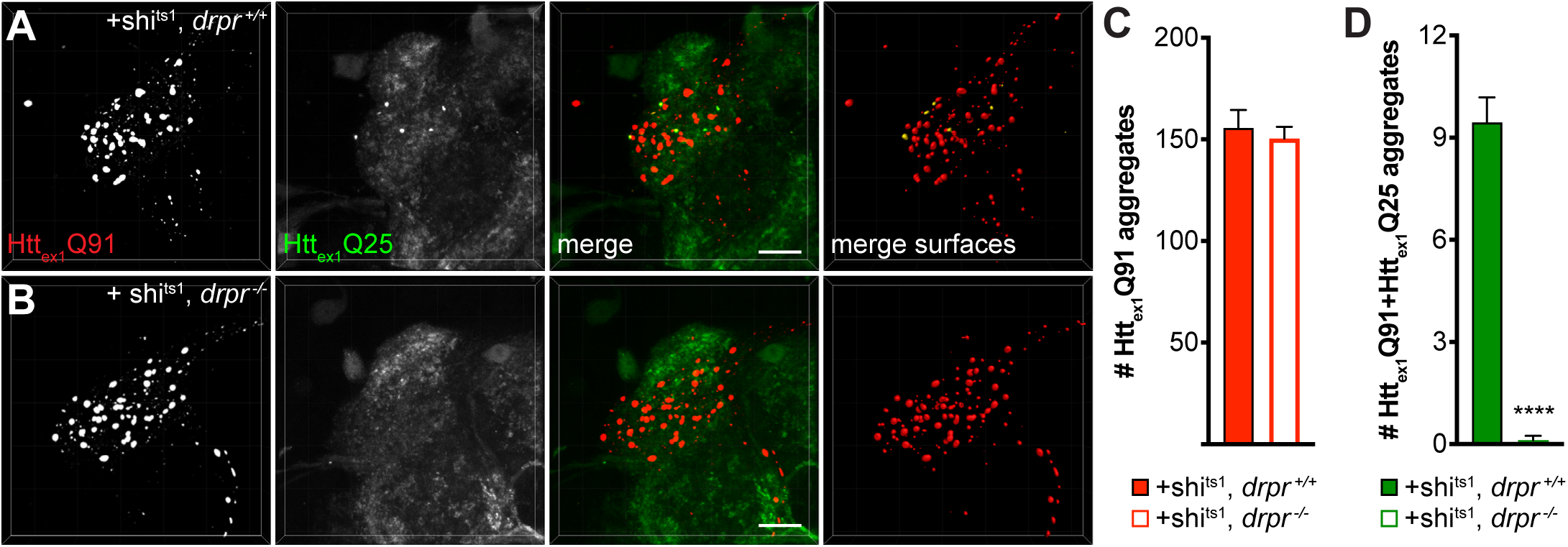
Draper is required for enhanced transfer of mHtt_ex1_ from shi^ts1^- expressing DA1 ORNs to GH146+ PNs. (A and B) Confocal stacks of DA1 glomeruli from 7 day-old *drpr ^+/+^* (A) or *drpr ^-/-^* (B) adult females co-expressing Htt_ex1_Q91-mCherry with shi^ts1^ in DA1 ORNs and Htt_ex1_Q25-GFP in GH146+ PNs. Adult flies were shifted from 18°C to 31°C upon eclosion. mCherry+ surfaces identified by semi-automated image segmentation are shown in the last panels, with Htt_ex1_Q91-only surfaces in red and Htt_ex1_Q91+Htt_ex1_Q25 surfaces in yellow. Scale bars = 10 μm. (C and D) Quantification of Htt_ex1_Q91 (C) and Htt_ex1_Q91+Htt_ex1_Q25 (D) aggregates for the same genotypes shown in (A and B). Data are shown as mean ± SEM; ****p < 0.0001 by Student’s t-test comparing *drpr ^+/+^* vs *drpr ^-/-^* flies.

**Figure 5 – figure supplement 2.**
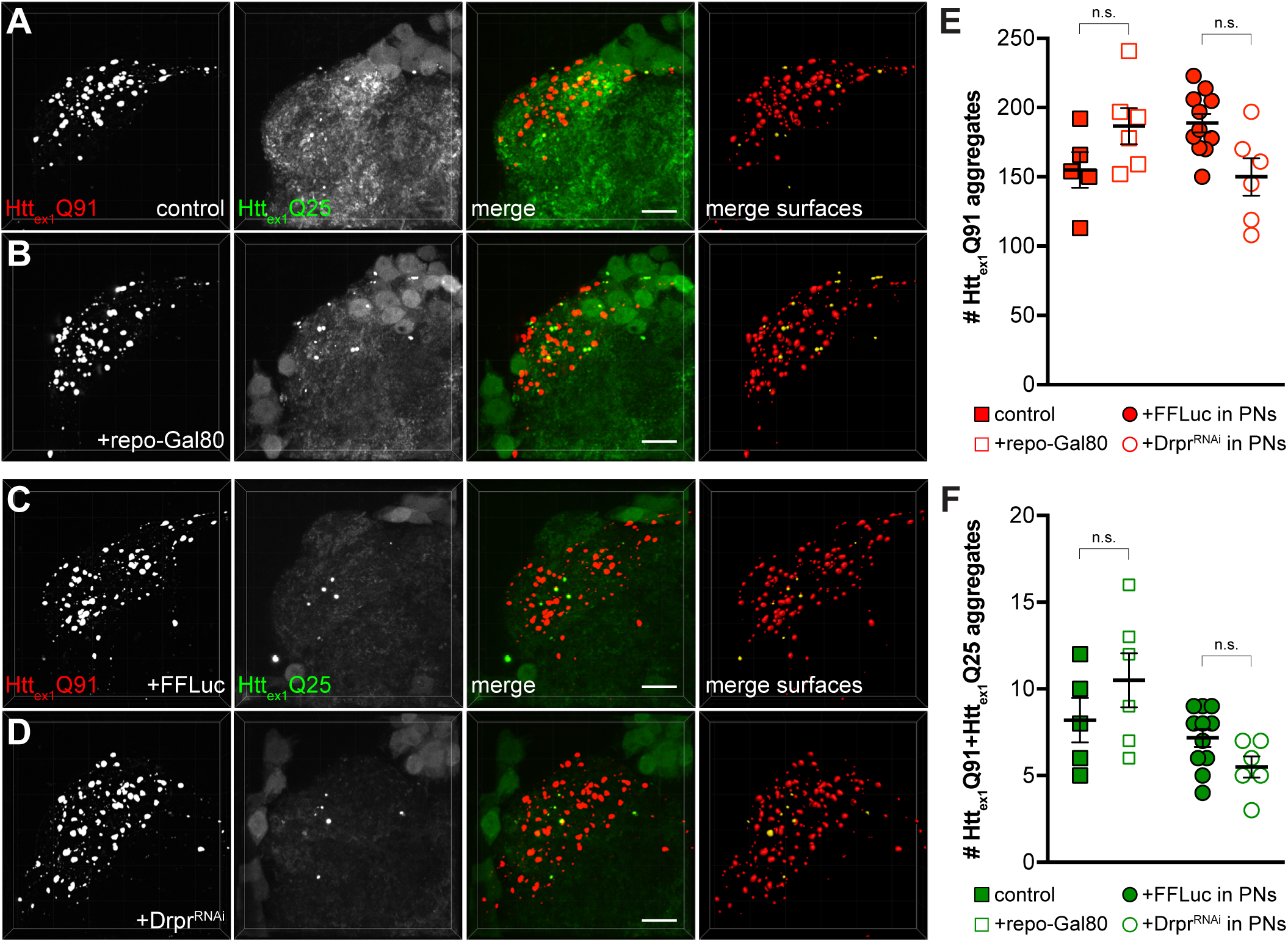
Gal80-mediated repression of Gal4 in glia or RNAi knockdown of drpr in PNs do not alter ORN-to-PN prion-like transfer of mHtt_ex1_ aggregates. (A and B) Confocal stacks of DA1 glomeruli from 10-14 day-old adult males expressing Htt_ex1_Q91-mCherry in DA1 ORNs and Htt_ex1_Q25-GFP in GH146+ PNs with (A) no additional transgenes (“control”) or (B) repo-Gal80 to inhibit Gal4-mediated expression of Htt_ex1_Q25-GFP in glia. (C and D) Confocal z-stacks of DA1 glomeruli from 14 day-old adult males expressing Htt_ex1_Q91-mCherry in DA1 ORNs and co-expressing Htt_ex1_Q25-GFP together with firefly luciferase (FFLuc; C) or dsRNA targeting draper (Drpr^RNAi^; D) in GH146+ PNs. In (A-D), mCherry+ surfaces identified by semi-automated image segmentation are shown in the last panels, with Htt_ex1_Q91-only surfaces in red and Htt_ex1_Q91+Htt_ex1_Q25 surfaces in yellow. Scale bars = 10 μm. (E and F) Quantification of Htt_ex1_Q91 (E) and Htt_ex1_Q91+Htt_ex1_Q25 (F) aggregates for genotypes shown in (A-D). Data are shown as mean ± SEM; n.s. = not significant by one-way ANOVA with Tukey’s multiple comparisons tests.

**Figure 5 – figure supplement 3.**
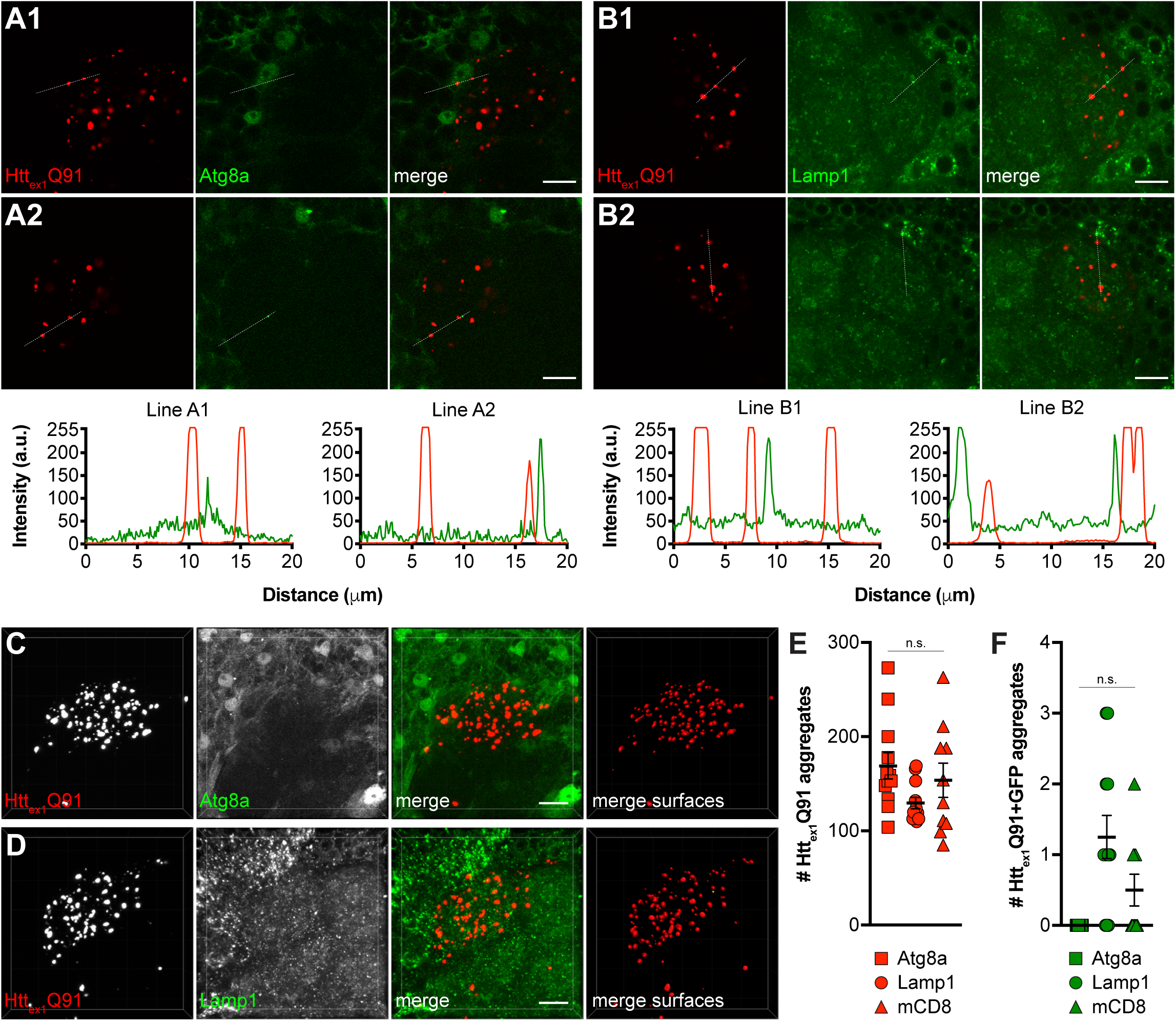
mHtt_ex1_ aggregates generated in ORNs do not co-localize with markers of lysosomes or autophagosomes in glia. (A1-2 and B1-2) Single confocal slices from DA1 glomeruli of 4-5 day-old adult males expressing Htt_ex1_Q91-mCherry in DA1 ORNs and either the autophagosomal marker Atg8a-GFP (A1 and A2) or the lysosomal marker GFP-Lamp1 (B1 and B2) in repo+ glia. Scale bars = 10 μm. Htt_ex1_Q91-mCherry (red) and Atg8a- or Lamp1-GFP (green) fluorescence intensity profiles for lines indicated in (A1-2 and B1-2) are shown below images; a.u. = arbitrary units. Lines were scanned from leftmost to rightmost point. (C and D) Confocal stacks of DA1 glomeruli from 4-5 day-old adult males expressing Htt_ex1_Q91-mCherry in DA1 ORNs and either Atg8a- GFP (C) or GFP-Lamp1 (D) in repo+ glia. mCherry+ surfaces identified by semi-automated image segmentation are shown in the last panels, with Htt_ex1_Q91-only surfaces in red and Htt_ex1_Q91+GFP surfaces in yellow. Scale bars = 10 μm. (E and F) Quantification of Htt_ex1_Q91 (E) and Htt_ex1_Q91+GFP (F) surfaces for the same genotypes as in (C and D) and for control animals expressing mCD8-GFP in glia. Data are shown as mean ± SEM; n.s. = not significant by one-way ANOVA with Tukey’s multiple comparisons test.

**Figure 7 - figure supplement 1.**
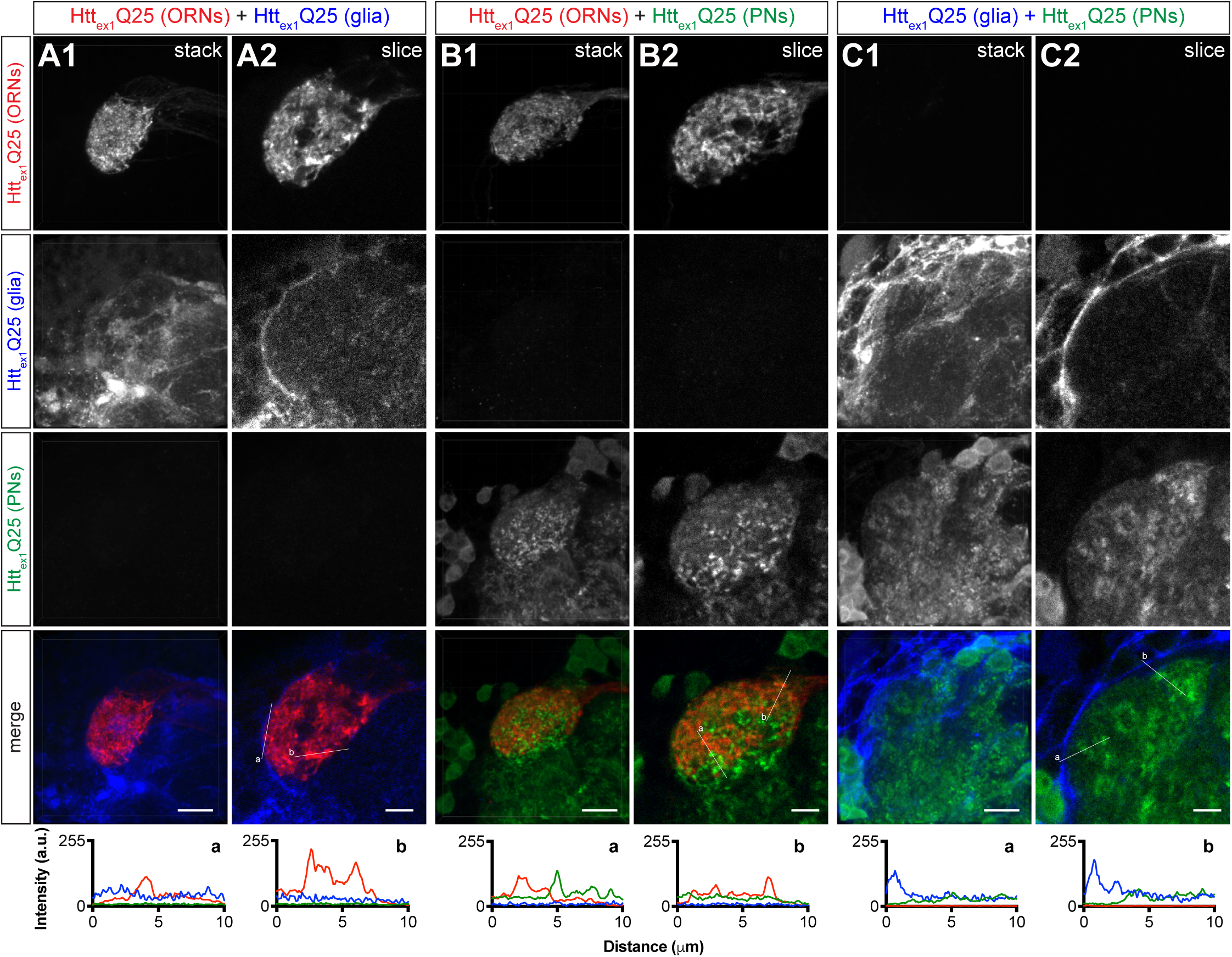
Controls for monitoring transmission of mHtt_ex1_ aggregates from presynaptic DA1 ORNs to postsynaptic PNs via a glial intermediate. **(A-C) Confocal z-stacks (A1,** B1, and C1) and single 0.35 μm z-slices (A2, B2, and C2) of DA1 glomeruli from 4-6 day-old adult females expressing (A1-2) Htt_ex1_Q25-mCherry in DA1 ORNs and Ht_tex1_Q25-3xHA in repo+ glia, (B1-2) Htt_ex1_Q25-mCherry in DA1 ORNs and Htt_ex1_Q25-YFP in GH146+ PNs, or (C1-2) Htt_ex1_Q25-3xHA in repo+ glia and Htt_ex1_Q25-YFP in GH146+ PNs. All brains were immunostained with antibodies against all three epitope tags (i.e., mCherry, YFP, and 3xHA) to show specific staining. Scale bars = 10 μm in (A1, B1, and C1) and 5 μm in (A2, B2, and C2). Htt_ex1_Q25-mCherry (red), Htt_ex1_Q25-3xHA (blue), and Htt_ex1_Q25-YFP (green) fluorescence intensity profiles for lines “a” and “b” shown in merged slices (A2, B2, and C2) are shown below images. Lines were scanned from leftmost to rightmost point.

**Video 1. Semi-automated quantification of mHtt_ex1_ and seeded wtHtt_ex1_ aggregates in the DA1 glomerulus**. Animation illustrating semi-automated approach for quantifying mHtt_ex1_ and wtHtt_ex1_ aggregates in brains expressing Htt_ex1_Q91-mCherry in DA1 ORNs and Htt_ex1_Q25-GFP in GH146+ PNs. Data shown in video is the same as in Fig. 1-figure supplement 2A1-2 and C1-7. Segmentation of raw high-magnification 3D confocal data (0:00) in the red channel (0:07) identified distinct Htt_ex1_Q91- mCherry surfaces (0:09), which were filtered for those that co-localize with high-intensity GFP signal to isolate the subpopulation associated with Htt_ex1_Q25-GFP puncta (0:13). Volumetric surfaces representing Htt_ex1_Q91 (*red*) and Htt_ex1_Q91+Htt_ex1_Q25 (*yellow*) aggregates (0:18) are shown for each data set analyzed by this method. The animation also illustrates segmentation of raw data in the green channel (0:28) to identify the brightest Htt_ex1_Q25-GFP objects in each data set (0:29). Overlap of mCherry+ and GFP+ surfaces, with GFP+ surfaces set at 50% transparency (0:30), highlights co-localization of smaller Htt_ex1_Q91 “seeds” surrounded by Htt_ex1_Q25 signal in these aggregates.

